# RCPedia: A global resource for studying and exploring retrocopies in diverse species

**DOI:** 10.1101/2023.12.20.572530

**Authors:** Helena B. Conceição, Rafael L. V. Mercuri, Matheus P. M. de Castro, Daniel T. Ohara, Gabriela D. A. Guardia, Pedro A F Galante

## Abstract

**Motivation:** Gene retrocopies, or processed pseudogenes, arise from the reverse transcription and genomic insertion of processed mRNA transcripts. These elements have significantly contributed to genetic diversity and novelties throughout the evolution of many species. However, the study of retrocopies has been challenging, owing to the absence of comprehensive, complete, and user-friendly databases for diverse species.

**Results:** Here, we introduce an improved version of RCPedia, an integrative database meticulously designed for the study of retrocopies. RCPedia offers an extensive catalog of retrocopies identified across 44 species, which includes 13 primates, 4 rodents, 6 chiropterans, 12 other mammals, 4 birds, turtle, lizard, frog, zebrafish, and drosophila. The database offers the most complete compilation of retrocopies per species, accompanied by detailed genomic annotations, expression data, and links to other data portals. Furthermore, RCPedia features a streamlined representation of data and an efficient querying system, establishing it as an invaluable tool for researchers in the fields of genomics, evolutionary biology and transposable elements. In summary, RCPedia aims to enhance the investigation of retrocopies and their pivotal roles in shaping the genomic landscapes of diverse species.

**Availability:** RCPedia is available at https://www.rcpediadb.org

## 1 INTRODUCTION

Gene retrocopies, or processed pseudogenes, are formed through the reverse transcription of mRNA, followed by genomic integration (Kaessmann et al., 2009). Opposed to their usual classification as processed pseudogenes, retrocopies have been recognized as functional genomic elements (Navarro and Galante, 2015; Casola and Betrán, 2017; Cheetham *et al.*, 2020). Their significance manifests in several aspects: (i) gene regulation, influencing the transcription of nearby or parental genes (Poliseno *et al.*, 2010); (ii) genetic innovation, with some retrocopies transcribed and translated into functional retrogenes distinct from parent genes (Burki and Kaessmann, 2004); (iii) contribution to genomic diversity, particularly through polymorphisms in populations (Schrider *et al.*, 2013); (iv) disease associations, including cancer, through gene function disruption or misregulation (Karreth *et al.*, 2015; Bim *et al.*, 2019). Thus, studying retrocopies is crucial for understanding the complex aspects of species evolution and innovation.

The advancement of genomics and related fields has been significantly accelerated by web tools and databases. These resources are indispensable for managing, analyzing, and interpreting extensive genomic datasets, facilitating groundbreaking discoveries. However, there is a noticeable scarcity of databases dedicated to retrocopies. Currently, only two such dedicated and active databases exist: RCPedia (Navarro and Galante, 2013) and RetrogeneDB (Kabza *et al.*, 2014). RCPedia was initially focused on human and primate retrocopies but was limited in species coverage. RetrogeneDB, while valuable, is based on a stringent identification pipeline resulting in a limited number of retrocopies per species. Additionally, expression data is restricted to few tissues, even in well-studied species like humans.

This improved version of RCPedia (https://www.rcpediadb.org) represents a significant advancement over its predecessor. Building on the original platform’s success, the new RCPedia version was built upon improved computational pipelines and up to date genomic references and gene annotations, offering an expanded and comprehensive catalog of retrocopies across 44 species, along with extensive RNA-Seq expression data. All features are available through a user-friendly web interface, enhancing accessibility and utility for the research community. This expansion and refinement of RCPedia are poised to substantially impact the study of retrocopies and their role in shaping the genomic landscape across diverse species.

## 2 DATA RETRIEVAL AND CURATION

### 2.1 Data sources

RCPedia includes data from 44 species, from human to drosophila (Supplementary Table 1). As with the previous version, the identification of retrocopies for each species is grounded in two key datasets: a well-assembled reference genome sequence and a set of known annotated genes. Figure 1A presents a schematic overview of the retrocopy identification process. A comprehensive description of our algorithm for retrocopy identification is provided in the supplementary material (and section below). The versions of the genomes and corresponding reference transcriptomes utilized are detailed in Supplementary Table 2. Retrocopy expression data are derived from publicly available RNA-seq datasets, the complete list of datasets used can be found in Supplementary Table 3.

**Figure 1.**
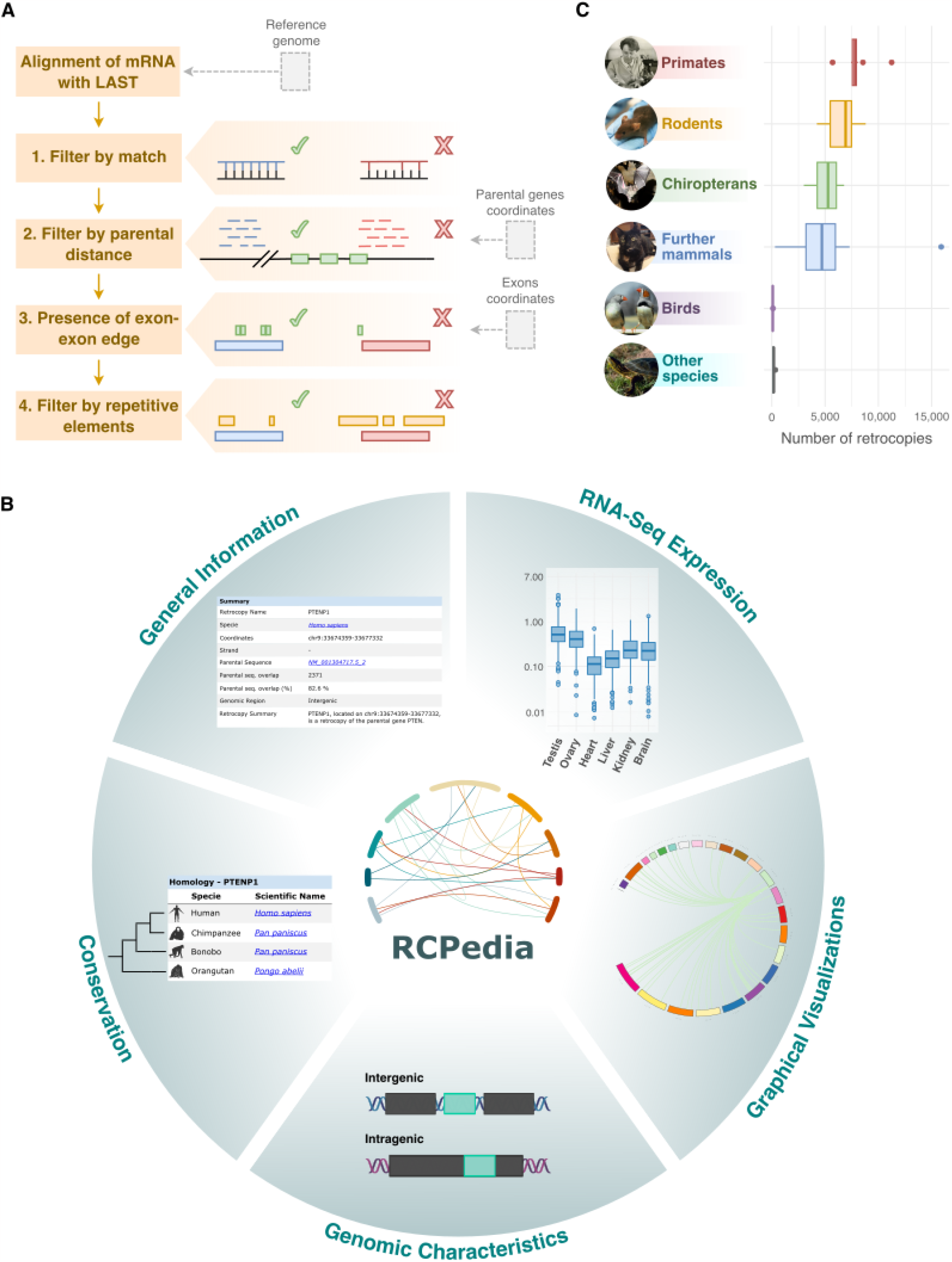
Overview of RCPedia’s Features and Data. A) The retrocopy identification workflow in RCPedia. This schematic delineates the algorithm’s main steps for identifying retrocopies, beginning with mRNA alignment and followed by sequential filtering based on match quality, parental gene distance, presence of exon-exon junctions, and exclusion of repetitive elements. B) The user interface and data presentation in RCPedia. Illustrated here is a sample of the comprehensive information available for each retrocopy and its parental gene, including summary statistics, genomic coordinates, cross-species homology, RNA-seq expression levels, and genomic context. C) Distribution of retrocopy numbers across different species groups. The box plots demonstrate the range and median values of retrocopies identified within groups of primates, rodents, chiropterans, other mammals, birds, and additional vertebrate species.

### 2.2 Identifying expressed and orthologs retrocopies

A crucial step for retrocopies to become functional is the acquisition of expression (Navarro and Galante, 2015; Carelli *et al.*, 2016). To provide the most comprehensive and accurate data on retrocopy expression, we reprocessed (or used pre-calculated expression data) from approximately 310 thousand samples across these 44 species. In humans and mice, we observed retrocopy expression in 308,406 samples spanning 73 tissues. For other species, expression was noted in six pre-selected tissues (brain, heart, liver, kidney, ovary, and testis) from a total of 1,580 samples. These tissues were specifically chosen due to their biological significance, the abundance of available RNA-seq data across many species, and their tendency for expressing retrocopies.

## 3 DATABASE IMPLEMENTATION

Similar to its previous version (Navarro and Galante, 2013), the RCPedia database is built on a relational database structure using MariaDB (https://mariadb.org/). The website was developed primarily in PHP (http://www.php.net), utilizing CakePHP (http://cakephp.org) as the framework for an efficient Model-View-Controller front-end. Genomes and gene annotations were processed using a combination of Perl (http://www.perl.org), Python, and in-house developed shell script algorithms.

For retrocopy identification, we refined our previously developed methods (Navarro and Galante, 2013, 2015). Briefly, coding transcripts from RefSeq were aligned against their respective reference genomes using Last (Kiełbasa *et al.*, 2011) with specified parameters. Our four-step strategy for identifying retrocopies involves: 1) Selecting alignments with at least 120 matched nucleotides;. 2) excluding alignments of messenger RNA in the original protein-coding gene loci or up to 200,000 bp of distance; 3) mapping exon-exon boundary positions from parental genes onto the alignments, retaining only those depicting an intronless region nearest the 3’ end of the messenger RNA; and 4) removing alignments that are composed in more than 40% of Transposable Elements (TEs) like LINEs and SINEs. The final retrocopy set was defined by selecting all remaining alignments and grouping those mapped to the same genomic locus (detailed in Supplementary Material).

## 4 DATABASE QUERY INTERFACE AND OUTPUT VISUALIZATION

### 4.1 The query system

RCPedia features both a direct and an advanced query system, designed for ease of use and speed (Supplementary Figure 1). Users can efficiently perform searches using commonly utilized identifiers. Queries can be made using the name of the parent gene (e.g., GAPDH), the retrocopy name (e.g., PTENP1), specific chromosomes (e.g., chr17), genomic positions (e.g., chr17:17000000-18000000), or across an entire chromosome. Additionally, the system supports queries based on other gene identifiers, such as ENSEMBL gene names (e.g., ENSG00000232230), and transcript identifiers, including Refseq IDs (e.g., NM_001281497.2), Supplementary Figure 2.

### 4.2 Results

RCPedia effectively presents information on retrocopies and parental genes for all 44 species analyzed. The database includes detailed annotation information for both genes and retrocopies, based on NCBI gene data. For humans, when a retrocopy is already named, we adhere to this annotation. Unannotated retrocopies are named following the pattern ‘parental gene name’ + P [number] (e.g., FTLP17). Additionally, we provide their genomic locations, sequences, and graphical visualizations through Circos plots. RCPedia also includes data on expression (measured in Transcripts per Million and the logarithm of this value) and the conservation of each retrocopy across species. This information is graphically represented in Figure 1B and Supplementary Figure 3. Figure 1C summarizes the number of retrocopies identified in each species group. Overall, Figure 1C shows that the number of retrocopies varies considerably across the groups. Primates and rodents exhibit higher and also variable numbers of retrocopies, while birds and other vertebrates (species) show fewer retrocopies with less variability. Interestingly, the presence of outliers in primates and rodents may suggest that certain species within these groups may possess unique evolutionary pressures or mechanisms influencing retrocopy numbers. Users can easily search for retrocopy numbers per species using RCPedia, as detailed in Supplementary Figure 4.

## 5 USING RCPedia

To demonstrate the utility of RCPedia, we focused on the retrocopies of the Glyceraldehyde-3-phosphate dehydrogenase (GAPDH) gene. GAPDH, a gene encoding a crucial enzyme in cellular metabolism, is involved in various cellular processes and is commonly used as a housekeeping gene due to its stable expression across tissues. However, it is also one of the most frequently retrocopied genes in mammals, a significant aspect often overlooked. RCPedia reveals 1,659 retrocopies of GAPDH across 29 species. Notably, rats, mice, and humans have 261, 100, and 53 GAPDH retrocopies, respectively, contributing to a total of 415 copies. Intriguingly, 789 of these retrocopies are conserved across other animals in our dataset, and 100% of species exhibit expression (> 0.1 TPM) in at least one of their GAPDH retrocopies in one or more samples. Specifically, in humans and mice, 52 and 96 retrocopies, respectively, are expressed (Supplementary Figure 5).

## 6 FUTURE PERSPECTIVES

In future updates of RCPedia, we aim to enrich the database with data that strengthens the evidence for retrocopy expression, shedding light on the regulatory mechanisms and further functional roles of these gene duplicates. By leveraging datasets from large-scale public initiatives, such as ENCODE, we aim to augment our database with Cap Analysis of Gene Expression (CAGE) for pinpointing transcription start sites of retrocopies in organisms like humans and mice. Additionally, we plan to integrate Assay for Transposase-Accessible Chromatin sequencing (ATAC-seq) to assess chromatin accessibility at retrocopy insertion sites, along with Chromatin Immunoprecipitation Sequencing (ChIP-Seq) to examine markers of transcriptional activity, including RNA Polymerase II occupancy and histone modifications associated with retrocopy expression. As the sequencing and annotation of genomes from more species improve, our efforts will continue in the identification and integration of retrocopies into RCPedia. Particularly for humans, we intend to expand our database to include RNA-Seq expression data from various pathological contexts, including cancer, to provide a more comprehensive resource for understanding retrocopy dynamics in disease.

## 7 CONCLUSION

RCPedia emerges as a well-organized, user-friendly resource featuring a streamlined graphical interface, focusing on the study of retrocopies in 44 species. This database significantly complements existing resources such as RetrogeneDB, Pseudogene.org, and retroFinder. It fills critical gaps by providing a dedicated platform for comprehensive retrocopy identification and enhances data integration, browsing, and visualization capabilities. We are confident that this updated and enhanced version of RCPedia will greatly facilitate more extensive and in-depth investigations of retrocopies across diverse species. This advancement is expected to deepen our understanding of retrocopies’ roles in genomic evolution, gene expression regulation, and their association with diseases.

## ACKNOWLEDGEMENTS

The authors would like to express their gratitude to Fabio C. P. Navarro and the members of Galante’s lab for their valuable suggestions.

