## Supplementary material for "RCPedia: A global resource for studying and exploring retrocopies in diverse species"

### Supplementary Figures

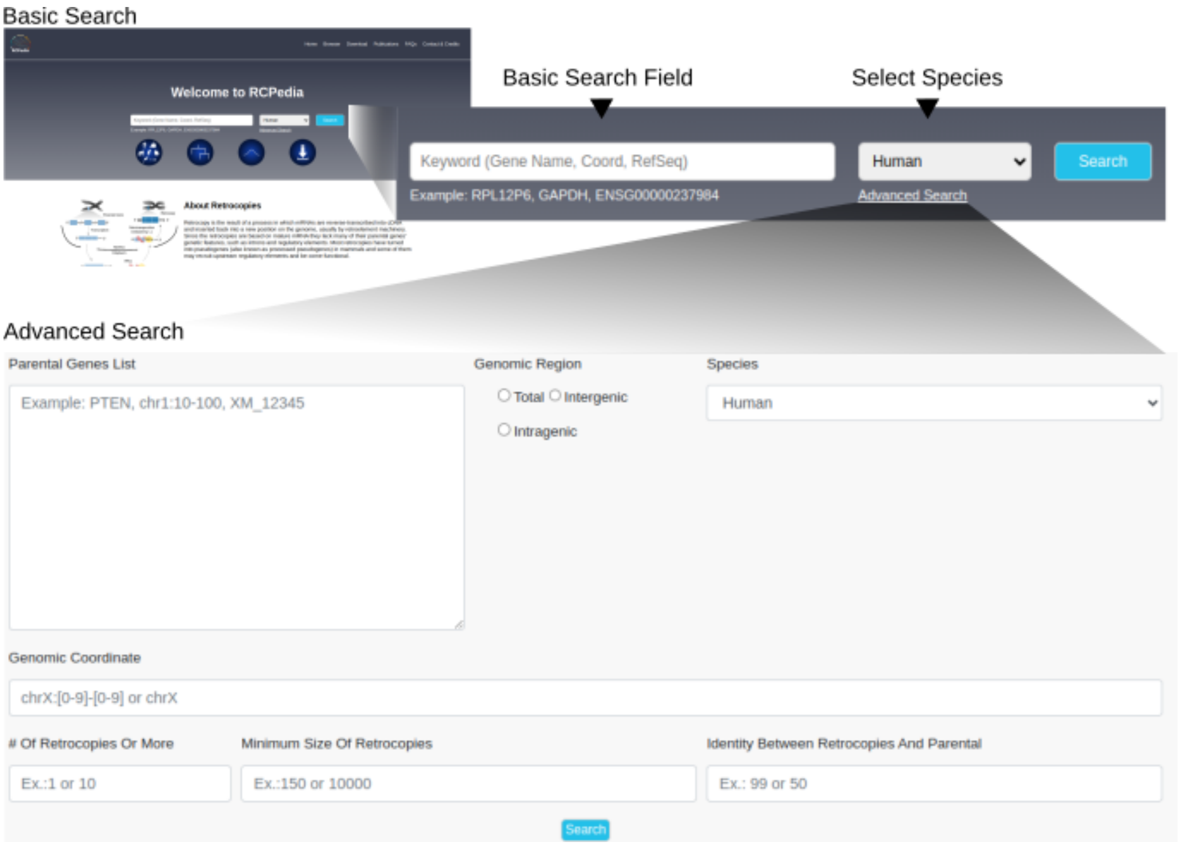

**Supplementary Figure 1.** RCPedia features both a direct and an advanced query system, designed for ease of use and speed.

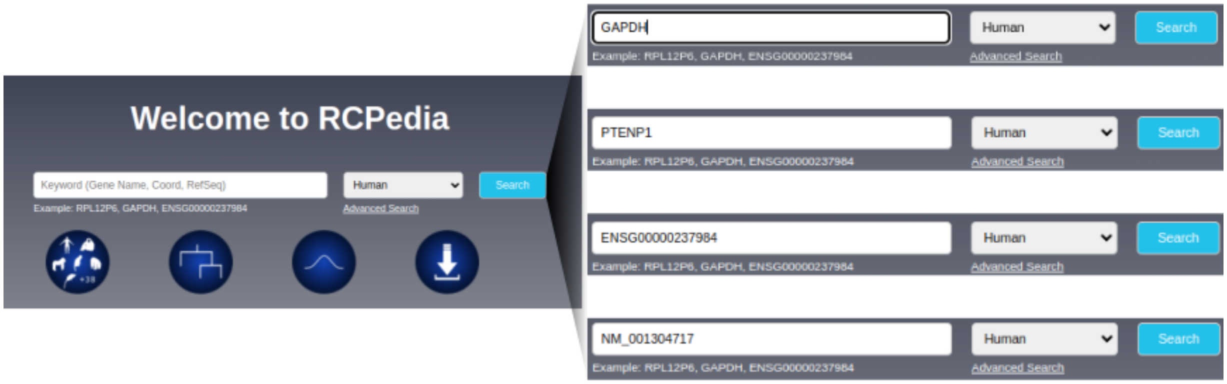


**Supplementary Figure 2. Users can efficiently perform searches using a variety of information**. Users can query using official gene names, including retrocopy names, ID from ENSEMBL and RefSeq database.

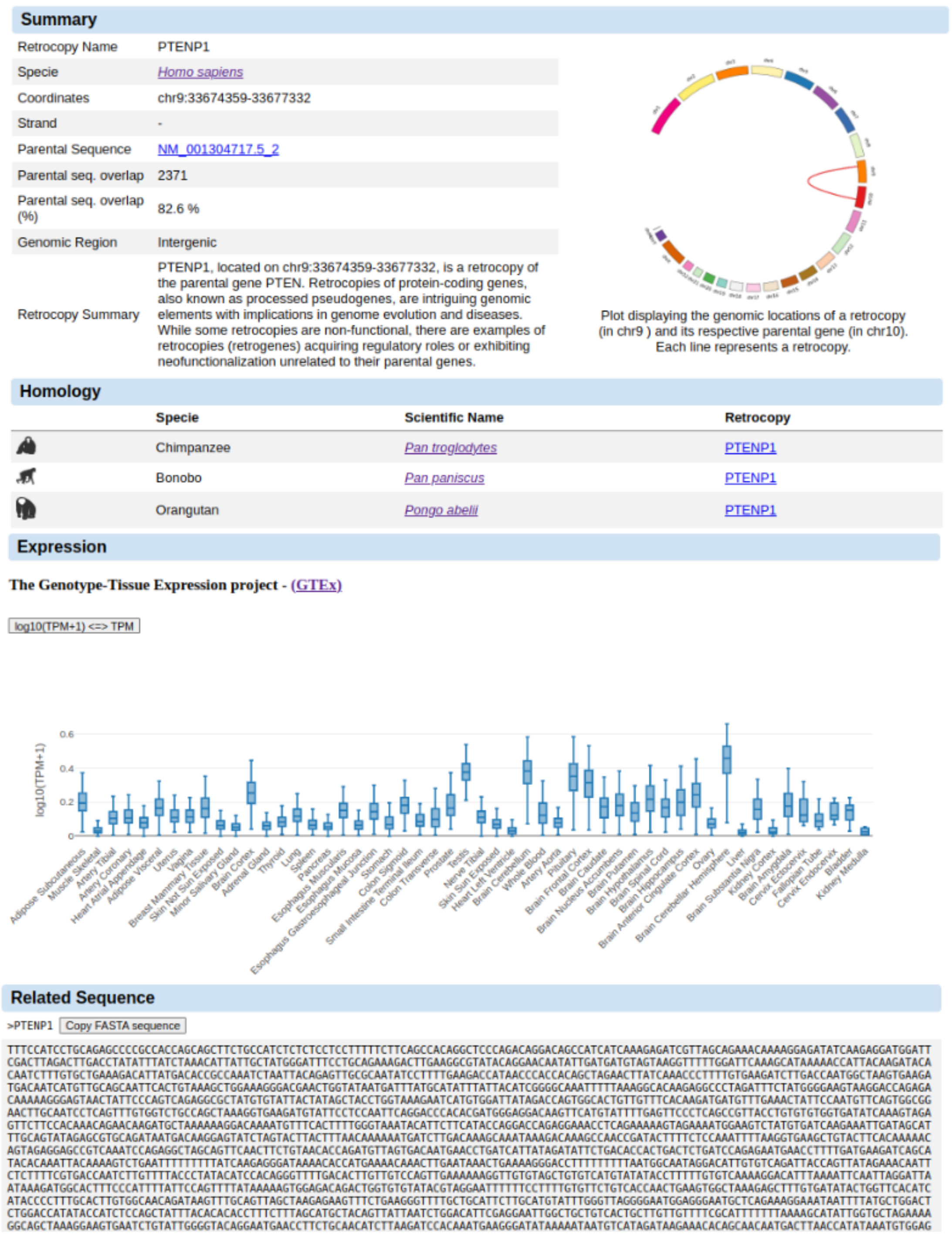


**Supplementary Figure 3. Comprehensive data on retrocopies and parental genes as presented by RCPedia.** The resource includes information on genomic locations and sequences, along with graphical visualizations such as Circos plots.

Additionally, RCPedia provides expression data (quantified in Transcripts per Million and logarithmic values) and details on the conservation of each retrocopy across different species.

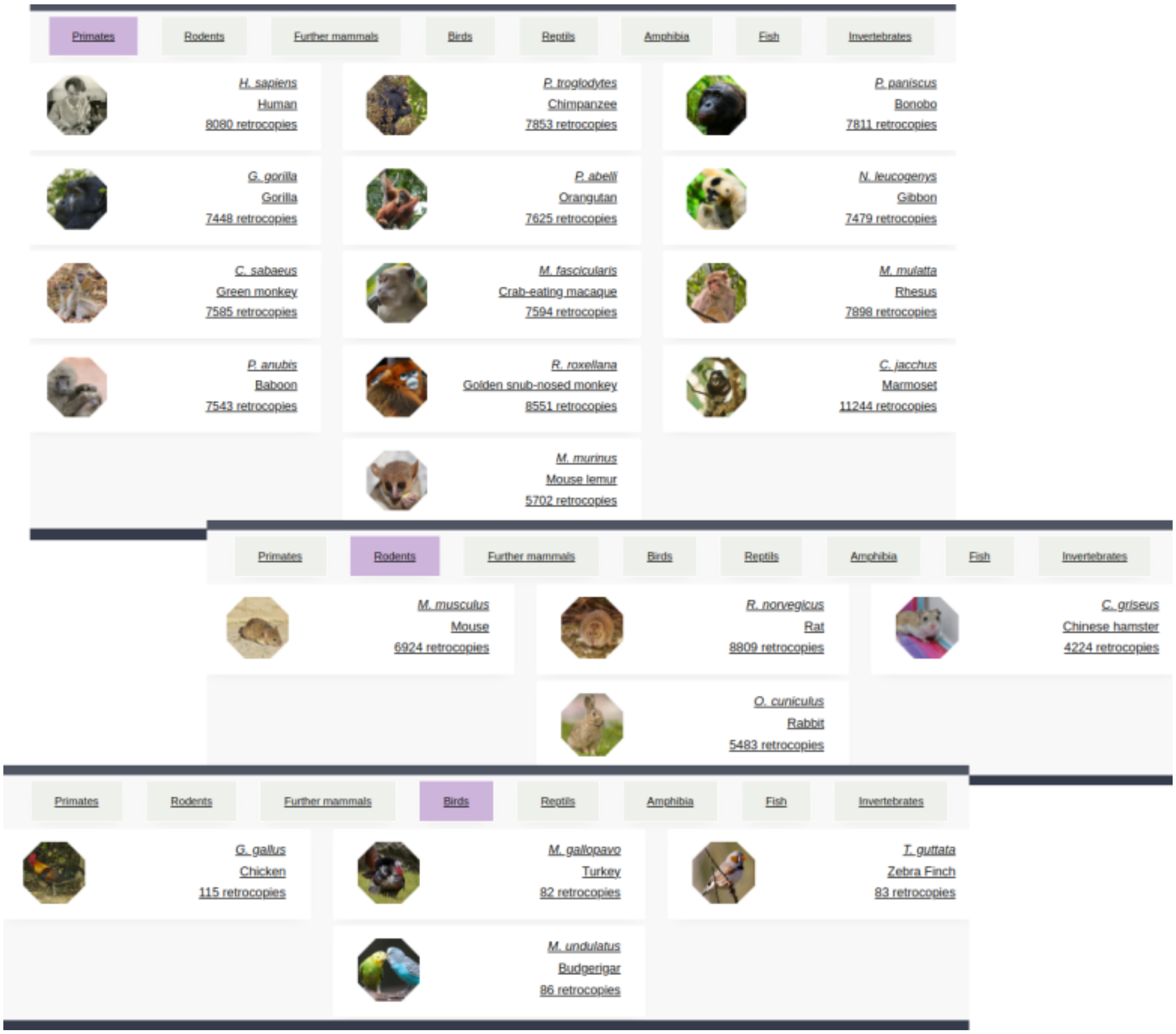


**Supplementary Figure 4. Simplified search for retrocopy numbers by species in RCPedia.** This figure provides a summary of the number of retrocopies for selected groups such as primates, rodents, and birds.

### Parental information

#### Summary

| | |
|---|---|
| Gene Name | Gapdh |
| Specie | *Rattus norvegicus* |
| Full Name | glyceraldehyde-3-phosphate dehydrogenase |
| Also known as | BARS-38\|Gapd |
| Coordinate | chr4:157676336-157680322 |
| Strand | - |
| Gene summary | This gene encodes a member of the glyceraldehyde-3-phosphate dehydrogenase protein family. A similar protein in human and mouse has been identified as a moonlighting protein based on its ability to perform mechanistically distinct functions. The encoded protein was originally identified as a key glycolytic enzyme that converts D-glyceraldehyde 3-phosphate (G3P) into 3-phospho-D-glyceroyl phosphate. Subsequent studies in human and mouse have assigned a variety of additional functions to the protein including nitrosylation of nuclear proteins. Many pseudogenes similar to this locus are found throughout the rat genome. [provided by RefSeq, Jan 2014] |

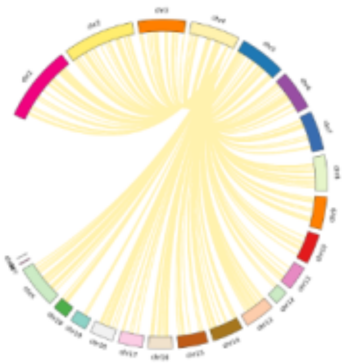

Plot displaying the genomic locations of a retrocopy and its respective parental gene (in chr4). Each line represents a retrocopy.

#### Retrocopy(s) from Gapdh

| Retroname | Coord | Strand | Genomic Region | ENSG | |
|---|---|---|---|---|---|
| GapdhP1 | chr1:105780971-105782338 | - | Intragenic | N/A | UCSC |
| GapdhP2 | chr1:107155915-107156995 | - | Intergenic | N/A | UCSC |
| GapdhP3 | chr1:137513948-137515169 | - | Intergenic | N/A | UCSC |
| GapdhP4 | chr1:167654597-167655479 | + | Intergenic | N/A | UCSC |

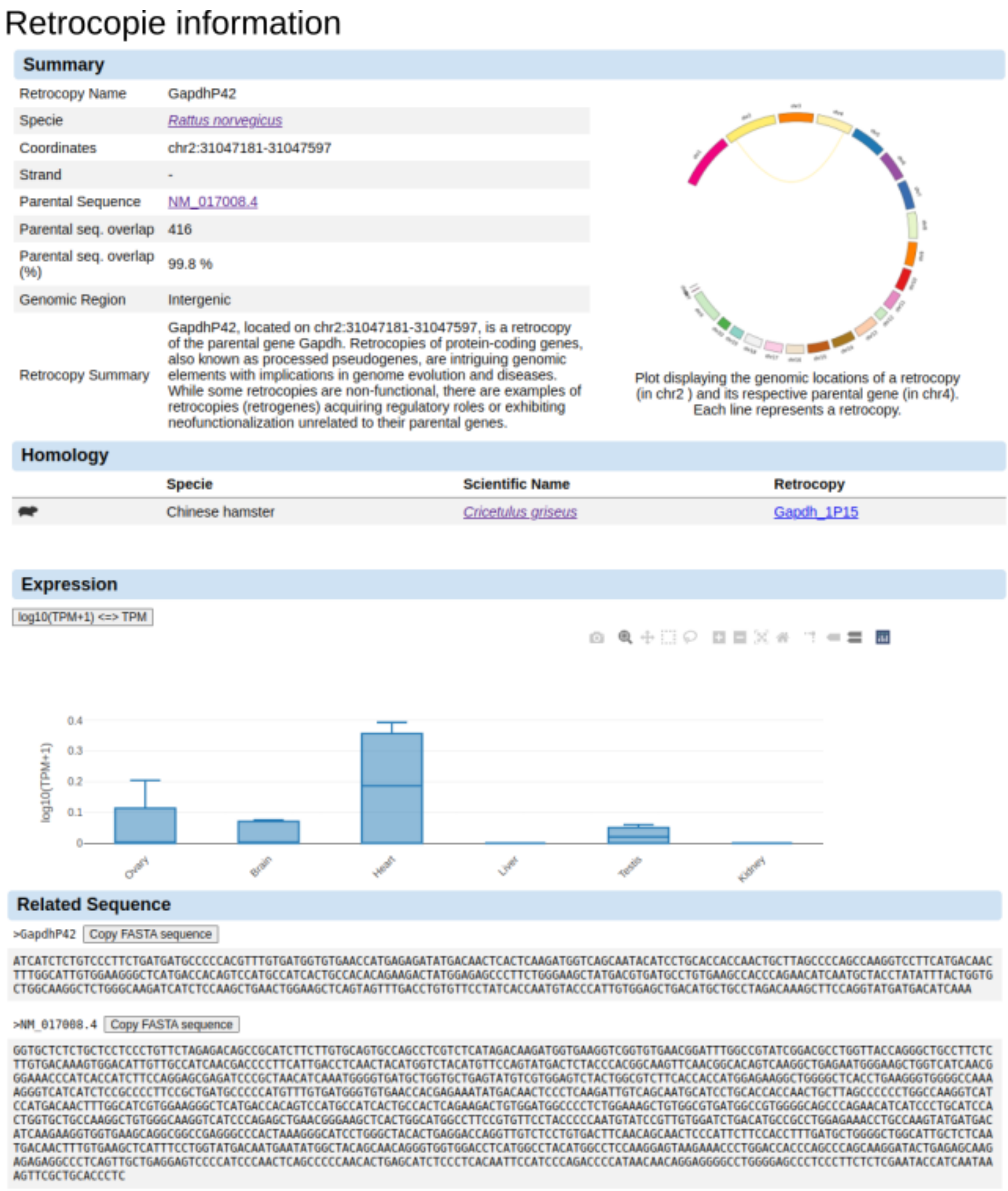


**Supplementary Figure 5. Gapdh and its retrocopies in rats.** This figure presents information from RCPedia on Gapdh (the parental gene) and its retrocopies. It includes gene annotation data and a Circos plot illustrating the genomic locations of Gapdh and its retrocopies.

### Supplementary Materials and Methods

**Retrocopies Identification**

Similar to the previous version of RCPedia, the retrocopy search begins with the alignment of messenger RNAs against the reference genome. Two major improvements were implemented: first, the sequence of messenger RNAs from coding genes (XM_ or NM_) is extracted from the genomic position annotated by RefSeq (in a gff file) using the gffread algorithm(Pertea and Pertea, n.d.). This step, avoids discrepancies between mRNA sequence and genomic annotation, an occasional inconsistency pointed out by RefSeq itself. Second, we transitioned from the BLAT aligner to the LAST aligner(Kiełbasa et al. 2011), executed with the parameter: lastal -D1000.

The pipeline incorporates a series of filters implemented through Python, Perl, and shell scripts, alongside bioinformatics tools like bedtools(Quinlan and Hall 2010). These filters include criteria such as match length, exon-exon boundary determination, alignment position, and repetitive element exclusion, resulting in a more accurate retrocopy identification.

Specifically: i) from the LAST output, we selected only alignments with a match exceeding 120 base pairs; ii) we selected only alignments with a distance greater than 200,000 base pairs from the origin gene of that messenger RNA, to remove possible chromosomal duplications; iii) we determined the exon-exon boundary position within the messenger RNA sequence of coding genes. We then checked the presence of the last, penultimate, or antepenultimate exon-exon boundary in the alignment, as the reverse transcriptase processes messenger RNA from the poly-A tail; iv) we excluded alignments composed mostly of repetitive elements (greater than or equal to 40%), based on RepeatMasker(“RepeatMasker Home Page” n.d.) annotation of repetitive elements, with the addition of simpleRepeats and windowMasker for non-mammals, .

In the case of multiple alignments from different messenger RNAs of the same gene, the alignment with the highest match is selected, and in the case of a tie, the one with the highest score (match/(match+mismatch)) is chosen. Additionally, the algorithm gathers continuous alignments originating from the same messenger RNA that are up to 6,000 base pairs apart in the genome, allowing for the presence of repetitive elements between shorter alignments.

To address challenges in non-mammalian species, a distance filter is applied, restricting retrocopy insertions from the same parental gene to every 500,000 base pairs, minimizing false positives without relying on pre-defined blacklists. Additionally, issues related to large gene families are handled by removing retrocopies overlapping 3 or more exons from annotated coding genes and by removing parental genes with more than 5 retrocopies that overlap with annotated genes from the same gene family. Lastly, candidates from distinct parental genes are selected based on best score, with ties broken randomly. Finally, manual analysis is conducted to retain occurrences of retrocopy insertions within older retrocopies. These improvements collectively enhance the precision and efficiency of retrocopy identification, particularly in non-mammalian species with distinct genomic characteristics.

**Retrocopies Homology**

To identify potential orthologous retrocopies, our approach involved retrieving the genomic sequence surrounding each retrocopy (3,000 base pairs upstream and downstream). We conducted pairwise alignment, employing the Lastz aligner(“Improved Pairwise Alignment of Genomic DNA” 2007), between each region and all corresponding regions of retrocopies (3,000 base pairs upstream and downstream) from other species. Filtering criteria were established, requiring alignment coverage exceeding 60% and identity surpassing 70% for comparisons within primates. For non-primate species or mixed species comparisons, a more permissive threshold of coverage above 50% and identity above 60% was applied, acknowledging the anticipated reduced conservation. Additionally, we verified if at least 60% of the retrocopy aligned with the "target" region of the other species in primate comparisons, or at least 50% in non-primate cases. In instances of multiple potential orthologs, preference was given to the alignment with the highest coverage and identity for each retrocopy. Regrettably, orthologs could not be identified for reptiles, amphibians, fish, and invertebrates (five species) due to the insufficient identity and coverage of the alignments.

**Retrocopies Expression based on RNA-Seq data**

Retrocopy expression quantification was conducted through the application of the Kallisto pseudo-alignment algorithm(Bray et al. 2016) in RNASeq experiments involving diverse tissues. Selected datasets from the NCBI SRA database were utilized for this purpose. The derived expression estimates underwent normalization, factoring in both the total number of mapped reads and transcript length. This normalization process yielded TPM (transcripts per million mapped reads) values, providing a standardized metric for retrocopy expression across various tissues.

Bray, Nicolas L., Harold Pimentel, Páll Melsted, and Lior Pachter. 2016. "Near-Optimal Probabilistic RNA-Seq Quantification." *Nature Biotechnology* 34 (5): 525–27.
"Improved Pairwise Alignment of Genomic DNA." 2007. https://search.proquest.com/openview/bc77cca0fb9390b44b9ef572fb574322/1?pq-origsite=gscholar&cbl=18750.
Kiełbasa, Szymon M., Raymond Wan, Kengo Sato, Paul Horton, and Martin C. Frith. 2011. "Adaptive Seeds Tame Genomic Sequence Comparison." *Genome Research* 21 (3): 487–93.
Pertea, G., and M. Pertea. n.d. "GFF Utilities: GffRead and GffCompare [version 1; Peer Review: 2." https://doi.org/10.12688/f1000research.23297.1.
Quinlan, Aaron R., and Ira M. Hall. 2010. "BEDTools: A Flexible Suite of Utilities for Comparing Genomic Features." *Bioinformatics* 26 (6): 841–42.
"RepeatMasker Home Page." n.d. Accessed December 20, 2023. http://www.repeatmasker.org.

**Supplementary Table 1. Species analyzed and present in the RCPedia database**

| Species (common name) | Species (scientific name) |
|---|---|
| Human | *Homo sapiens* |
| Chimpanzee | *Pan troglodytes* |
| Bonobo | *Pan paniscus* |
| Gorilla | *Gorilla gorilla gorilla* |
| Orangutan | *Pongo pygmaeus abelii* |
| Gibbon | *Nomascus leucogenys* |
| Green monkey | *Chlorocebus sabaeus* |
| Crab-eating macaque | *Macaca fascicularis* |
| Rhesus | *Macaca mulatta* |
| Baboon (anubis) | *Papio anubis* |
| Golden snub-nosed monkey | *Rhinopithecus roxellana* |
| Marmoset | *Callithrix jacchus* |
| Mouse lemur | *Microcebus murinus* |
| Mouse 2020 | *Mus musculus* |
| Rat | *Rattus norvegicus* |
| Chinese hamster | *Cricetulus griseus* |
| Rabbit | *Oryctolagus cuniculus* |
| Pig | *Sus scrofa* |
| Cow | *Bos taurus* |
| Sheep | *Ovis aries* |
| Dolphin | *Tursiops truncatus* |
| Horse | *Equus caballus* |
| Dog | *Canis lupus familiaris* |
| Panda | *Ailuropoda melanoleuca* |
| Cat | *Felis catus* |
| Pale spear-nosed bat | *Phyllostomus discolor* |
| Greater horseshoe bat | *Rhinolophus ferrumequinum* |

| | |
|---|---|
| Velvety free-tailed bat | *Molossus molossus* |
| Egyptian rousette | *Rousettus aegyptiacus* |
| Kuhl's pipistrelle | *Pipistrellus kuhlii* |
| Greater mouse-eared bat | *Myotis myotis* |
| Sloth | *Choloepus hoffmanni* |
| Tasmanian devil | *Sarcophilus harrisii* |
| Opossum | *Monodelphis domestica* |
| Platypus | *Ornithorhynchus anatinus* |
| Chicken | *Gallus gallus* |
| Turkey | *Meleagris gallopavo* |
| Zebra finch | *Taeniopygia guttata* |
| Budgerigar | *Melopsittacus undulatus* |
| Painted turtle | *Chrysemys picta bellii* |
| Lizard | *Anolis carolinensis* |
| X tropicalis | *Xenopus tropicalis* |
| Zebrafish | *Danio rerio* |
| D melanogaster | *Drosophila melanogaster* |

**Supplementary Table 2. Reference genomes used in RCPedia**

| Species | Reference Genomes | Assembly ID |
|---|---|---|
| Human | hg38 | GCF_000001405.39 |
| Chimpanzee | panTro6 | GCF_002880755.1 |
| Bonobo | panPan3 | GCF_013052645.1 |
| Gorilla | gorGor6 | GCF_008122165.1 |
| Orangutan | ponAbe3 | GCF_002880775.1 |
| Gibbon | Asia_NLE_v1 | GCF_006542625.1 |
| Green monkey | chlSab2 | GCF_000409795.2 |
| Crab-eating macaque | macFas5 | GCF_000364345.1 |
| Rhesus | rheMac10 | GCF_003339765.1 |
| Baboon | Panubis1.0 | GCF_008728515.1 |
| Golden snub-nosed monkey | ASM756505v1 | GCF_007565055.1 |
| Marmoset | calJac4 | GCF_009663435.1 |
| Mouse lemur | Mmur_3.0 | GCF_000165445.2 |
| Mouse 2020 | mm39 | GCF_000001635.27 |
| Rat | rn6 | GCF_000001895.5 |
| Chinese hamster | CriGri-PICRH-1.0 | GCF_003668045.3 |
| Rabbit | oryCun2 | GCF_000003625.3 |
| Pig | susScr11 | GCF_000003025.6 |
| Cow | bosTau9 | GCF_002263795.1 |
| Sheep | Oar_rambouillet_v1.0 | GCF_002742125.1 |
| Dolphin | mTurTru1.mat.Y | GCF_011762595.1 |
| Horse | equCab3 | GCF_002863925.1 |
| Dog | ROS_Cfam_1.0 | GCF_014441545.1 |
| Panda | ASM200744v2 | GCF_002007445.1 |
| Cat | felCat9 | GCF_000181335.3 |
| Sloth | mChoDid1.pri | GCF_015220235.1 |
| Tasmanian Devil | mSarHar1.11 | GCF_902635505.1 |
| Opossum | monDom5 | GCF_000002295.2 |

| | | |
|---|---|---|
| Platypus | mOrnAna1.p.v1 | GCF_004115215.1 |
| Chicken | galGal6 | GCF_000002315.6 |
| Turkey | Turkey_5.1 | GCF_000146605.3 |
| Zebra Finch | bTaeGut2.pat.W.v2 | GCF_008822105.2 |
| Budgerigar | bMelUnd1.mat.Z | GCF_012275295.1 |
| Painted Turtle | Chrysemys_picta_BioNano-3.0.4 | GCF_000241765.4 |
| Lizard | anoCar2 | GCF_000090745.1 |
| Frog | xenTro10 | GCF_000004195.4 |
| Zebrafish | danRer11 | GCF_000002035.6 |
| Drosophila | dm6 | GCF_000001215.4 |
| Pale spear-nosed bat | mPhyDis1.pri.v3 | GCF_004126475.2 |
| Greater horseshoe bat | mRhiFer1_v1.p | GCF_004115265.1 |
| Pale spear-nosed bat | HLphyDis3 | GCA_014049915.1 |
| Greater horseshoe bat | HLrhiFer5 | GCA_014108255.1 |
| Velvety free-tailed bat | HLmolMol2 | GCF_014108415.1 |
| Egyptian rousette | HLrouAeg4 | GCF_014176215.1 |
| Kuhl's pipistrelle | HLpipKuh2 | GCF_014108245.1 |
| Greater mouse-eared bat | HLmyoMyo6 | GCF_014108235.1 |

**Supplementary Table 3. RNA Sequencing Data Utilized for Retrocopy Quantification in RCPedia.** The raw FASTQ files were downloaded, reprocessed, and the expressions of retrocopies (and their parental genes) were quantified. These data are available in RCPedia.

| Species Tax ID | Sample ID* | Tissues |
|---|---|---|
| 208526 | SRS454178 | testis |
| 208526 | SRS454179 | ovary |
| 208526 | SRS454180 | heart |
| 208526 | SRS454181 | heart |
| 61622 | SRR9417645 | Heart |
| 60711 | SRR5275322 | Brain |
| 60711 | SRR5275323 | Brain |
| 60711 | SRR5275325 | Brain |
| 60711 | SRR5275328 | Brain |
| 60711 | SRR5275329 | Brain |
| 60711 | SRR5275331 | Brain |
| 60711 | SRR6010875 | Brain |
| 60711 | SRR6010878 | Brain |
| 60711 | SRR6010883 | Brain |
| 60711 | SRR6010884 | Brain |
| 60711 | SRR6010889 | Brain |
| 60711 | SRR6010891 | Brain |
| 60711 | SRR6010901 | Brain |
| 60711 | SRR6010916 | Brain |
| 60711 | SRR6010920 | Brain |
| 60711 | SRR8062405 | Kidney |
| 60711 | SRR8062406 | Kidney |
| 60711 | SRR8062407 | Kidney |
| 60711 | SRR8475000 | Kidney |
| 60711 | SRR8475001 | Kidney |

| | | |
|---|---|---|
| 60711 | SRR8475002 | Kidney |
| 60711 | SRR8475003 | Kidney |
| 60711 | SRR8475004 | Kidney |
| 60711 | SRR8475005 | Kidney |
| 59729 | SRR13358559 | Brain |
| 59729 | SRR13358570 | Brain |
| 59729 | SRR13358585 | Brain |
| 59729 | SRR18111875 | Brain |
| 59729 | SRR18111876 | Brain |
| 59729 | SRR18111877 | Brain |
| 59729 | SRR18111878 | Brain |
| 59729 | SRR18111879 | Brain |
| 59729 | SRR18111880 | Brain |
| 59729 | SRR18111881 | Brain |
| 59729 | SRR18111882 | Brain |
| 59729 | SRR18111883 | Brain |
| 59729 | SRR18111884 | Brain |
| 59729 | SRR18111885 | Brain |
| 59729 | SRR18111886 | Brain |
| 30608 | SRR1758989 | Brain |
| 30608 | SRR1758990 | Brain |
| 30608 | SRR1758991 | Brain |
| 30608 | SRR1758992 | Brain |
| 30608 | SRR1758993 | Brain |
| 30608 | SRR1758995 | Kidney |
| 30608 | SRR1758996 | Kidney |
| 30608 | SRR1758997 | Liver |
| 30608 | SRR7704801 | Testis |
| 28377 | SRR11050685 | Liver |
| 28377 | SRR11050686 | Testis |

| | | |
|---|---|---|
| 28377 | SRR11050687 | Testis |
| 28377 | SRR579556 | Brain |
| 28377 | SRR579557 | Kidney |
| 28377 | SRR579558 | Heart |
| 28377 | SRR8700054 | Kidney |
| 28377 | SRR8700055 | Testis |
| 28377 | SRR8700057 | Liver |
| 28377 | SRR8700058 | Heart |
| 28377 | SRR8700061 | Brain |
| 28377 | SRR8700062 | Heart |
| 28377 | SRR8700063 | Kidney |
| 28377 | SRS282426 | heart |
| 28377 | SRS282427 | ovary |
| 28377 | SRS282428 | brain |
| 28377 | SRS283111 | liver |
| 28377 | SRS366869 | brain |
| 28377 | SRS366870 | kidney |
| 28377 | SRS366871 | heart |
| 27675 | SRR10066841 | Brain |
| 27675 | SRR10066842 | Liver |
| 27675 | SRR10066844 | Kidney |
| 27675 | SRR11580490 | Brain |
| 27675 | SRR11580492 | Liver |
| 27675 | SRR11580494 | Kidney |
| 13616 | ERS1090504 | liver |
| 13616 | ERS1090519 | liver |
| 13616 | SRR12341176 | Kidney |
| 13616 | SRR12341177 | Kidney |
| 13616 | SRR12341179 | Kidney |
| 13616 | SRR12341180 | Kidney |

| | | |
|---|---|---|
| 13616 | SRR12341182 | Kidney |
| 13616 | SRR12341183 | Kidney |
| 13616 | SRR13142088 | Liver |
| 13616 | SRR13142089 | Liver |
| 13616 | SRR13142090 | Liver |
| 13616 | SRR13142094 | Brain |
| 13616 | SRR13142095 | Brain |
| 13616 | SRR13142096 | Brain |
| 13616 | SRR13142100 | Testis |
| 13616 | SRR13142101 | Testis |
| 13616 | SRR13142102 | Testis |
| 13616 | SRR6206899 | Brain |
| 13616 | SRR6206904 | Heart |
| 13616 | SRR6206909 | Kidney |
| 13616 | SRR6206914 | Liver |
| 13616 | SRR6206928 | Testis |
| 13616 | SRS304760 | brain |
| 13616 | SRS304761 | brain |
| 13616 | SRS304762 | liver |
| 13616 | SRS335256 | brain |
| 13616 | SRS335257 | testis |
| 13616 | SRS335259 | heart |
| 13616 | SRS335260 | liver |
| 13616 | SRS335263 | ovary |
| 13616 | SRS335265 | kidney |
| 13616 | SRS429621 | ovary |
| 13616 | SRS429819 | testis |
| 13616 | SRS448463 | heart |
| 13616 | SRS448464 | kidney |
| 13616 | SRS448465 | liver |

| | | |
|---|---|---|
| 13616 | SRS449815 | brain |
| 13616 | SRS449818 | liver |
| 13616 | SRS449820 | testis |
| 10116 | ERR3384586 | Brain |
| 10116 | ERR3384592 | Brain |
| 10116 | ERR3384600 | Brain |
| 10116 | ERR3384602 | Brain |
| 10116 | ERR3384652 | Brain |
| 10116 | ERR3384711 | Brain |
| 10116 | ERR3384728 | Brain |
| 10116 | ERR3384733 | Brain |
| 10116 | ERR3384735 | Brain |
| 10116 | ERR3384794 | Brain |
| 10116 | ERR3384800 | Brain |
| 10116 | ERR3384801 | Brain |
| 10116 | ERR3384807 | Brain |
| 10116 | ERR3384808 | Brain |
| 10116 | ERR3384816 | Brain |
| 10116 | ERR3417900 | Testis |
| 10116 | ERR3417906 | Testis |
| 10116 | ERR3417950 | Testis |
| 10116 | ERR3995074 | Heart |
| 10116 | ERR3995075 | Heart |
| 10116 | ERR3995076 | Heart |
| 10116 | ERR3995077 | Heart |
| 10116 | ERR3995078 | Heart |
| 10116 | ERR3995079 | Heart |
| 10116 | ERR3995080 | Heart |
| 10116 | ERR3995081 | Heart |
| 10116 | ERR3995082 | Heart |

| | | |
|---|---|---|
| 10116 | ERR3995083 | Heart |
| 10116 | ERR3995084 | Heart |
| 10116 | ERR3995085 | Heart |
| 10116 | ERR3995086 | Heart |
| 10116 | ERR3995087 | Heart |
| 10116 | ERR4881963 | Liver |
| 10116 | ERR4881965 | Liver |
| 10116 | ERR4881966 | Liver |
| 10116 | ERR4881972 | Liver |
| 10116 | ERR4881977 | Liver |
| 10116 | ERR4881979 | Liver |
| 10116 | ERR4881980 | Liver |
| 10116 | ERR4881981 | Liver |
| 10116 | ERR4881984 | Liver |
| 10116 | ERR4881986 | Liver |
| 10116 | ERR4881987 | Liver |
| 10116 | ERR4881988 | Liver |
| 10116 | ERR4881989 | Liver |
| 10116 | ERR4881991 | Liver |
| 10116 | ERR4881992 | Liver |
| 10116 | ERR4881993 | Heart |
| 10116 | ERR5924829 | Kidney |
| 10116 | ERR5924830 | Kidney |
| 10116 | ERR5924831 | Kidney |
| 10116 | ERR5924832 | Kidney |
| 10116 | ERR5924833 | Kidney |
| 10116 | ERR5924834 | Kidney |
| 10116 | ERR5924835 | Kidney |
| 10116 | ERR5924836 | Kidney |
| 10116 | ERR5924839 | Kidney |

| | | |
|---|---|---|
| 10116 | ERS523413 | brain |
| 10116 | ERS523414 | heart |
| 10116 | ERS523415 | kidney |
| 10116 | ERS523416 | liver |
| 10116 | ERS523419 | ovary |
| 10116 | ERS523422 | testis |
| 10116 | SRR3657387 | Ovary |
| 10116 | SRR3657388 | Ovary |
| 10116 | SRR5767265 | Kidney |
| 10116 | SRR5767266 | Kidney |
| 10116 | SRR5767267 | Kidney |
| 10116 | SRR5767268 | Kidney |
| 10116 | SRR5767269 | Kidney |
| 10116 | SRR5767270 | Kidney |
| 10116 | SRR922252 | Testis |
| 10116 | SRR922253 | Testis |
| 10116 | SRR922254 | Testis |
| 10116 | SRR922255 | Testis |
| 10116 | SRR922256 | Testis |
| 10116 | SRR922257 | Testis |
| 10116 | SRR922258 | Testis |
| 10116 | SRR922259 | Testis |
| 10116 | SRS1496490 | ovary |
| 10116 | SRS1496492 | ovary |
| 10116 | SRS369727 | brain |
| 10116 | SRS369729 | heart |
| 10116 | SRS369731 | liver |
| 10116 | SRS369735 | testis |
| 10116 | SRS369736 | brain |
| 10116 | SRS369738 | heart |

| | | |
|---|---|---|
| 10116 | SRS369740 | liver |
| 10116 | SRS369744 | testis |
| 10116 | SRS369745 | brain |
| 10116 | SRS369747 | heart |
| 10116 | SRS369748 | kidney |
| 10116 | SRS369749 | liver |
| 10116 | SRS369753 | testis |
| 10116 | SRS380729 | liver |
| 10116 | SRS380730 | liver |
| 10116 | SRS380731 | liver |
| 10116 | SRS380779 | kidney |
| 10116 | SRS380780 | kidney |
| 10116 | SRS380781 | kidney |
| 10116 | SRS380824 | brain |
| 10116 | SRS380825 | brain |
| 10116 | SRS380826 | brain |
| 10029 | ERR4319912 | Ovary |
| 10029 | ERR4319915 | Ovary |
| 10029 | ERR4319916 | Ovary |
| 10029 | ERR4319917 | Ovary |
| 10029 | ERR4319918 | Ovary |
| 10029 | ERR4319922 | Ovary |
| 10029 | ERR4319923 | Ovary |
| 10029 | ERR4319924 | Ovary |
| 10029 | ERR4319925 | Ovary |
| 10029 | ERR4319930 | Ovary |
| 10029 | ERR4319931 | Ovary |
| 10029 | ERR4319932 | Ovary |
| 10029 | ERR4319933 | Ovary |
| 10029 | ERR4319935 | Ovary |

| | | |
|---|---|---|
| 10029 | ERR4319936 | Ovary |
| 9986 | ERR3417971 | Testis |
| 9986 | ERR3417973 | Testis |
| 9986 | ERR3417980 | Testis |
| 9986 | ERS1090463 | liver |
| 9986 | ERS1090464 | liver |
| 9986 | ERS1090465 | liver |
| 9986 | SRR11939337 | Kidney |
| 9986 | SRR11939344 | Kidney |
| 9986 | SRR11939353 | Kidney |
| 9986 | SRR12012193 | Brain |
| 9986 | SRR12012194 | Brain |
| 9986 | SRR12012195 | Brain |
| 9986 | SRR12012196 | Brain |
| 9986 | SRR12012197 | Brain |
| 9986 | SRR12012198 | Brain |
| 9986 | SRR12012199 | Brain |
| 9986 | SRR12012200 | Brain |
| 9986 | SRR12012201 | Brain |
| 9986 | SRR12012202 | Brain |
| 9986 | SRR12012203 | Brain |
| 9986 | SRR12012204 | Brain |
| 9986 | SRR12012205 | Brain |
| 9986 | SRR12012206 | Brain |
| 9986 | SRR12012207 | Brain |
| 9986 | SRR12012208 | Brain |
| 9986 | SRR12012209 | Brain |
| 9986 | SRR12012210 | Brain |
| 9986 | SRR12012211 | Brain |
| 9986 | SRR12012212 | Brain |

| | | |
|---|---|---|
| 9986 | SRR12227967 | Testis |
| 9986 | SRR12227968 | Testis |
| 9986 | SRR12227969 | Testis |
| 9986 | SRR12227970 | Testis |
| 9986 | SRR12227971 | Testis |
| 9986 | SRR12227972 | Testis |
| 9986 | SRR12227973 | Testis |
| 9986 | SRR12227974 | Testis |
| 9986 | SRR12227983 | Testis |
| 9986 | SRR12227994 | Testis |
| 9986 | SRR12228005 | Testis |
| 9986 | SRR12228016 | Testis |
| 9986 | SRR12228021 | Testis |
| 9986 | SRR12228022 | Testis |
| 9986 | SRR12228023 | Testis |
| 9986 | SRR12228024 | Testis |
| 9986 | SRR17443340 | Heart |
| 9986 | SRR17443341 | Heart |
| 9986 | SRR17443362 | Heart |
| 9986 | SRR17443366 | Heart |
| 9986 | SRR17443367 | Heart |
| 9986 | SRR17443368 | Heart |
| 9986 | SRR17443373 | Heart |
| 9986 | SRR17443374 | Heart |
| 9986 | SRR17443375 | Heart |
| 9986 | SRR17443384 | Heart |
| 9986 | SRR17443387 | Heart |
| 9986 | SRR17443394 | Heart |
| 9986 | SRR17443395 | Heart |
| 9986 | SRR17443396 | Heart |

| | | |
|---|---|---|
| 9986 | SRR17443397 | Heart |
| 9986 | SRR17443398 | Heart |
| 9986 | SRR17443407 | Heart |
| 9986 | SRR17443412 | Heart |
| 9986 | SRR17443420 | Heart |
| 9986 | SRR17443421 | Heart |
| 9986 | SRR17443453 | Heart |
| 9986 | SRR17443454 | Heart |
| 9986 | SRR1786022 | Kidney |
| 9986 | SRR1786023 | Kidney |
| 9986 | SRR1786025 | Kidney |
| 9986 | SRR1786026 | Kidney |
| 9986 | SRR1789327 | Kidney |
| 9986 | SRR1789329 | Kidney |
| 9986 | SRR1789331 | Kidney |
| 9986 | SRR1789332 | Kidney |
| 9986 | SRR18329937 | Kidney |
| 9986 | SRR18329938 | Kidney |
| 9986 | SRR18329939 | Kidney |
| 9986 | SRR21082900 | Liver |
| 9986 | SRR21082901 | Liver |
| 9986 | SRR21082902 | Liver |
| 9986 | SRR21082903 | Liver |
| 9986 | SRR21082904 | Liver |
| 9986 | SRR21082905 | Liver |
| 9986 | SRR21082906 | Liver |
| 9986 | SRR21082907 | Liver |
| 9986 | SRR21082908 | Liver |
| 9986 | SRR21082909 | Liver |
| 9986 | SRR21082910 | Liver |

| | | |
|---|---|---|
| 9986 | SRR21082911 | Liver |
| 9986 | SRR21082912 | Liver |
| 9986 | SRR21082913 | Liver |
| 9986 | SRR21082914 | Liver |
| 9986 | SRR6206905 | Kidney |
| 9986 | SRR6206925 | Testis |
| 9986 | SRS072785 | ovary |
| 9986 | SRS072793 | liver |
| 9986 | SRS072798 | testis |
| 9986 | SRS072799 | heart |
| 9986 | SRS072802 | brain |
| 9986 | SRS072808 | kidney |
| 9986 | SRS380725 | liver |
| 9986 | SRS380726 | liver |
| 9986 | SRS380773 | kidney |
| 9986 | SRS380774 | kidney |
| 9986 | SRS380818 | brain |
| 9986 | SRS380819 | brain |
| 9986 | SRS836129 | kidney |
| 9986 | SRS836137 | kidney |
| 9986 | SRS836139 | kidney |
| 9986 | SRS836140 | kidney |
| 9986 | SRS836141 | heart |
| 9986 | SRS836142 | heart |
| 9986 | SRS836143 | liver |
| 9986 | SRS836144 | liver |
| 9986 | SRS836145 | heart |
| 9986 | SRS836147 | heart |
| 9986 | SRS836149 | liver |
| 9986 | SRS836150 | liver |

| | | |
|---|---|---|
| 9986 | SRS836178 | kidney |
| 9986 | SRS836179 | heart |
| 9986 | SRS836180 | kidney |
| 9986 | SRS836181 | kidney |
| 9986 | SRS836182 | heart |
| 9986 | SRS836183 | kidney |
| 9986 | SRS836184 | heart |
| 9986 | SRS836185 | heart |
| 9986 | SRS836186 | heart |
| 9986 | SRS836187 | heart |
| 9986 | SRS836188 | heart |
| 9986 | SRS836189 | heart |
| 9986 | SRS836190 | liver |
| 9986 | SRS836191 | liver |
| 9986 | SRS836192 | liver |
| 9986 | SRS836193 | liver |
| 9940 | ERR2074333 | Kidney |
| 9940 | ERR2074343 | Heart |
| 9940 | ERR2074344 | Heart |
| 9940 | ERR2074345 | Heart |
| 9940 | ERR2074346 | Heart |
| 9940 | ERR2074347 | Heart |
| 9940 | ERR2074348 | Heart |
| 9940 | ERR2074349 | Heart |
| 9940 | ERR2074350 | Heart |
| 9940 | ERR2074351 | Heart |
| 9940 | ERR2074352 | Heart |
| 9940 | ERR2074353 | Heart |
| 9940 | ERR2074354 | Heart |
| 9940 | ERR2074355 | Heart |

| | | |
|---|---|---|
| 9940 | ERR2074356 | Heart |
| 9940 | ERR2074357 | Heart |
| 9940 | ERR2074421 | Kidney |
| 9940 | ERR2074422 | Kidney |
| 9940 | ERR2074423 | Kidney |
| 9940 | ERR2074424 | Kidney |
| 9940 | ERR2074425 | Kidney |
| 9940 | ERR2074426 | Kidney |
| 9940 | ERR2075072 | Kidney |
| 9940 | ERR2075073 | Kidney |
| 9940 | ERR2075074 | Kidney |
| 9940 | ERR2075075 | Kidney |
| 9940 | ERR9627677 | Kidney |
| 9940 | ERR9627678 | Kidney |
| 9940 | ERR9627679 | Kidney |
| 9940 | ERR9627680 | Kidney |
| 9940 | ERS1869179 | ovary |
| 9940 | ERS1869181 | ovary |
| 9940 | ERS1869182 | ovary |
| 9940 | ERS2107387 | testis |
| 9940 | ERS444234 | ovary |
| 9940 | ERS444254 | ovary |
| 9940 | ERS444283 | brain |
| 9940 | ERS444285 | testis |
| 9940 | ERS444300 | liver |
| 9940 | ERS444304 | liver |
| 9940 | SRR15931175 | Testis |
| 9940 | SRR15931176 | Testis |
| 9940 | SRR15931177 | Testis |
| 9940 | SRR15931178 | Testis |

| | | |
|---|---|---|
| 9940 | SRR17030653 | Brain |
| 9940 | SRR17030654 | Brain |
| 9940 | SRR17030655 | Brain |
| 9940 | SRR17030656 | Brain |
| 9940 | SRR17030657 | Brain |
| 9940 | SRR17030658 | Brain |
| 9940 | SRR17030659 | Brain |
| 9940 | SRR17030660 | Brain |
| 9940 | SRR17030661 | Brain |
| 9940 | SRR17030662 | Brain |
| 9940 | SRR17030663 | Brain |
| 9940 | SRR17030664 | Brain |
| 9940 | SRR17299400 | Liver |
| 9940 | SRR17299401 | Liver |
| 9940 | SRR17299402 | Liver |
| 9940 | SRR17299403 | Liver |
| 9940 | SRR17299404 | Liver |
| 9940 | SRR17299405 | Liver |
| 9940 | SRR17299406 | Liver |
| 9940 | SRR17299407 | Liver |
| 9940 | SRR17299408 | Liver |
| 9940 | SRR17299409 | Liver |
| 9940 | SRR17299410 | Liver |
| 9940 | SRR17299411 | Liver |
| 9940 | SRR17299412 | Liver |
| 9940 | SRR17299413 | Liver |
| 9940 | SRR17299414 | Liver |
| 9940 | SRR20852782 | Testis |
| 9940 | SRR20852783 | Testis |
| 9940 | SRR20852784 | Testis |

| | | |
|---|---|---|
| 9940 | SRR20852785 | Testis |
| 9940 | SRR20852786 | Testis |
| 9940 | SRR20852787 | Testis |
| 9940 | SRR20852788 | Testis |
| 9940 | SRR20852789 | Testis |
| 9940 | SRR20852790 | Testis |
| 9940 | SRR20852791 | Testis |
| 9940 | SRR21021023 | Testis |
| 9940 | SRS1254769 | kidney |
| 9940 | SRS1254770 | kidney |
| 9940 | SRS1254771 | kidney |
| 9940 | SRS1254773 | liver |
| 9940 | SRS1254774 | liver |
| 9940 | SRS1254775 | heart |
| 9940 | SRS1254776 | heart |
| 9940 | SRS1254777 | heart |
| 9940 | SRS173290 | heart |
| 9940 | SRS173291 | heart |
| 9940 | SRS173295 | heart |
| 9940 | SRS173296 | heart |
| 9940 | SRS173297 | heart |
| 9940 | SRS173298 | heart |
| 9940 | SRS173299 | heart |
| 9940 | SRS1930467 | heart |
| 9940 | SRS1930470 | heart |
| 9940 | SRS1930473 | heart |
| 9940 | SRS1930528 | kidney |
| 9940 | SRS1930529 | kidney |
| 9940 | SRS1930530 | kidney |
| 9940 | SRS1930543 | liver |

| | | |
|---|---|---|
| 9940 | SRS1930545 | liver |
| 9940 | SRS1930547 | liver |
| 9940 | SRS2590160 | testis |
| 9940 | SRS2590682 | testis |
| 9940 | SRS2689185 | ovary |
| 9940 | SRS2689186 | ovary |
| 9940 | SRS2689187 | ovary |
| 9940 | SRS2689188 | ovary |
| 9940 | SRS2689189 | ovary |
| 9940 | SRS2689190 | ovary |
| 9940 | SRS2689191 | ovary |
| 9940 | SRS2689192 | ovary |
| 9940 | SRS2951364 | liver |
| 9940 | SRS2951365 | liver |
| 9940 | SRS2951366 | liver |
| 9940 | SRS2951367 | liver |
| 9940 | SRS2951369 | liver |
| 9940 | SRS2951370 | liver |
| 9940 | SRS2951371 | liver |
| 9940 | SRS590749 | liver |
| 9940 | SRS590750 | kidney |
| 9940 | SRS590751 | ovary |
| 9940 | SRS590752 | brain |
| 9940 | SRS590753 | heart |
| 9940 | SRS775844 | ovary |
| 9940 | SRS922843 | testis |
| 9940 | SRS922844 | testis |
| 9940 | SRS922845 | testis |
| 9940 | SRS922846 | testis |
| 9940 | SRS922847 | testis |

| | | |
|---|---|---|
| 9940 | SRS922848 | testis |
| 9940 | SRS922849 | testis |
| 9940 | SRS922850 | testis |
| 9940 | SRS922851 | testis |
| 9940 | SRS922852 | testis |
| 9940 | SRS922854 | testis |
| 9913 | SRR10031495 | Kidney |
| 9913 | SRR10031501 | Kidney |
| 9913 | SRR10031502 | Kidney |
| 9913 | SRR10736432 | Kidney |
| 9913 | SRR10736433 | Kidney |
| 9913 | SRR10736434 | Kidney |
| 9913 | SRR12108691 | Brain |
| 9913 | SRR12108692 | Brain |
| 9913 | SRR12108693 | Brain |
| 9913 | SRR12108694 | Brain |
| 9913 | SRR12108695 | Brain |
| 9913 | SRR12108696 | Brain |
| 9913 | SRR12108697 | Brain |
| 9913 | SRR12108698 | Brain |
| 9913 | SRR12108699 | Brain |
| 9913 | SRR12108700 | Brain |
| 9913 | SRR12108701 | Brain |
| 9913 | SRR12108702 | Brain |
| 9913 | SRR12108703 | Brain |
| 9913 | SRR12108704 | Brain |
| 9913 | SRR12108705 | Brain |
| 9913 | SRR12170551 | Heart |
| 9913 | SRR12170552 | Heart |
| 9913 | SRR12170553 | Heart |

| | | |
|---|---|---|
| 9913 | SRR12170554 | Heart |
| 9913 | SRR12170555 | Heart |
| 9913 | SRR12170556 | Heart |
| 9913 | SRR13204086 | Liver |
| 9913 | SRR13204087 | Liver |
| 9913 | SRR13204088 | Liver |
| 9913 | SRR13204089 | Liver |
| 9913 | SRR13204090 | Liver |
| 9913 | SRR13204091 | Liver |
| 9913 | SRR13204092 | Liver |
| 9913 | SRR13204093 | Liver |
| 9913 | SRR13204094 | Liver |
| 9913 | SRR13204095 | Liver |
| 9913 | SRR13204096 | Liver |
| 9913 | SRR13204097 | Liver |
| 9913 | SRR13204098 | Liver |
| 9913 | SRR13204099 | Liver |
| 9913 | SRR13204100 | Liver |
| 9913 | SRR17724234 | Testis |
| 9913 | SRR17724235 | Testis |
| 9913 | SRR17724237 | Testis |
| 9913 | SRR17724239 | Testis |
| 9913 | SRR17724240 | Testis |
| 9913 | SRR17724241 | Testis |
| 9913 | SRR17724244 | Testis |
| 9913 | SRR17724246 | Testis |
| 9913 | SRR17724247 | Testis |
| 9913 | SRR17724248 | Testis |
| 9913 | SRR17724254 | Testis |
| 9913 | SRR17724255 | Testis |

| | | |
|---|---|---|
| 9913 | SRR17724259 | Testis |
| 9913 | SRR17724260 | Testis |
| 9913 | SRR17724261 | Testis |
| 9913 | SRR19745551 | Kidney |
| 9913 | SRR19745554 | Kidney |
| 9913 | SRR19745555 | Kidney |
| 9913 | SRR6760949 | Heart |
| 9913 | SRR6760950 | Heart |
| 9913 | SRR6760951 | Heart |
| 9913 | SRR6760952 | Heart |
| 9913 | SRS1098356 | ovary |
| 9913 | SRS1254778 | kidney |
| 9913 | SRS1254779 | kidney |
| 9913 | SRS1254780 | kidney |
| 9913 | SRS1254784 | heart |
| 9913 | SRS1254785 | heart |
| 9913 | SRS1254786 | heart |
| 9913 | SRS369781 | brain |
| 9913 | SRS369783 | heart |
| 9913 | SRS369784 | kidney |
| 9913 | SRS369789 | testis |
| 9913 | SRS369790 | brain |
| 9913 | SRS369792 | heart |
| 9913 | SRS369793 | kidney |
| 9913 | SRS369798 | testis |
| 9913 | SRS369799 | brain |
| 9913 | SRS369801 | heart |
| 9913 | SRS369802 | kidney |
| 9913 | SRS369807 | testis |
| 9913 | SRS380743 | kidney |

| | | |
|---|---|---|
| 9913 | SRS380744 | kidney |
| 9913 | SRS380788 | brain |
| 9913 | SRS380789 | brain |
| 9913 | SRS485384 | testis |
| 9913 | SRS486117 | testis |
| 9913 | SRS625424 | heart |
| 9913 | SRS625430 | ovary |
| 9913 | SRS718130 | liver |
| 9913 | SRS718135 | liver |
| 9913 | SRS718142 | liver |
| 9913 | SRS718143 | liver |
| 9913 | SRS718144 | liver |
| 9913 | SRS718145 | liver |
| 9913 | SRS718146 | liver |
| 9913 | SRS718147 | liver |
| 9913 | SRS718148 | liver |
| 9913 | SRS718149 | liver |
| 9913 | SRS718150 | liver |
| 9913 | SRS718151 | liver |
| 9913 | SRS718152 | liver |
| 9913 | SRS718221 | liver |
| 9913 | SRS718223 | liver |
| 9913 | SRS734278 | testis |
| 9823 | ERS682351 | heart |
| 9823 | ERS682352 | kidney |
| 9823 | ERS762245 | heart |
| 9823 | ERS762246 | heart |
| 9823 | ERS762247 | heart |
| 9823 | ERS762248 | heart |
| 9823 | ERS762249 | heart |

| | | |
|---|---|---|
| 9823 | ERS762250 | heart |
| 9823 | ERS762251 | heart |
| 9823 | ERS762252 | heart |
| 9823 | SRR14193613 | Testis |
| 9823 | SRR14193614 | Testis |
| 9823 | SRR14193615 | Testis |
| 9823 | SRR14193616 | Testis |
| 9823 | SRR14193617 | Testis |
| 9823 | SRR14193618 | Testis |
| 9823 | SRR14193619 | Testis |
| 9823 | SRR14193620 | Testis |
| 9823 | SRR14193621 | Testis |
| 9823 | SRR14193622 | Testis |
| 9823 | SRR14193623 | Testis |
| 9823 | SRR14193624 | Testis |
| 9823 | SRR14241019 | Liver |
| 9823 | SRR14241020 | Liver |
| 9823 | SRR14241021 | Liver |
| 9823 | SRR14241022 | Liver |
| 9823 | SRR14241023 | Liver |
| 9823 | SRR14241024 | Liver |
| 9823 | SRR14241025 | Liver |
| 9823 | SRR14241026 | Liver |
| 9823 | SRR14241027 | Liver |
| 9823 | SRR14241028 | Liver |
| 9823 | SRR14241029 | Liver |
| 9823 | SRR14241030 | Liver |
| 9823 | SRR14241031 | Liver |
| 9823 | SRR14241032 | Liver |
| 9823 | SRR14583710 | Kidney |

| | | |
|---|---|---|
| 9823 | SRR14583722 | Kidney |
| 9823 | SRR14583728 | Kidney |
| 9823 | SRR14583732 | Kidney |
| 9823 | SRR14583735 | Kidney |
| 9823 | SRR14583757 | Kidney |
| 9823 | SRR15902208 | Brain |
| 9823 | SRR15902211 | Brain |
| 9823 | SRR15902213 | Brain |
| 9823 | SRR15902215 | Brain |
| 9823 | SRR15902216 | Brain |
| 9823 | SRR15902217 | Brain |
| 9823 | SRR15902220 | Brain |
| 9823 | SRR15902225 | Brain |
| 9823 | SRR15902227 | Brain |
| 9823 | SRR15902228 | Brain |
| 9823 | SRR15902229 | Brain |
| 9823 | SRR15902230 | Brain |
| 9823 | SRR15902231 | Brain |
| 9823 | SRR15902232 | Brain |
| 9823 | SRR15902236 | Brain |
| 9823 | SRR18494201 | Kidney |
| 9823 | SRR18494202 | Kidney |
| 9823 | SRR18494203 | Kidney |
| 9823 | SRR18494204 | Kidney |
| 9823 | SRR18494205 | Kidney |
| 9823 | SRR18494206 | Kidney |
| 9823 | SRR18494207 | Kidney |
| 9823 | SRR18494208 | Kidney |
| 9823 | SRR18494210 | Kidney |
| 9823 | SRR20073469 | Testis |

| | | |
|---|---|---|
| 9823 | SRR20073472 | Testis |
| 9823 | SRR20073473 | Testis |
| 9823 | SRR21023588 | Heart |
| 9823 | SRR21023596 | Heart |
| 9823 | SRR21023597 | Heart |
| 9823 | SRR21023598 | Heart |
| 9823 | SRR7630959 | Liver |
| 9823 | SRR9627715 | Heart |
| 9823 | SRR9627716 | Heart |
| 9823 | SRR9627717 | Heart |
| 9823 | SRR9627718 | Heart |
| 9823 | SRR9627719 | Heart |
| 9823 | SRR9627720 | Heart |
| 9823 | SRR9627721 | Heart |
| 9823 | SRR9627722 | Heart |
| 9823 | SRR9627723 | Heart |
| 9823 | SRR9627724 | Heart |
| 9823 | SRR9627725 | Heart |
| 9823 | SRS1094779 | ovary |
| 9823 | SRS1094780 | ovary |
| 9823 | SRS1094781 | ovary |
| 9823 | SRS1094782 | ovary |
| 9823 | SRS1094783 | ovary |
| 9823 | SRS1094784 | ovary |
| 9823 | SRS1100807 | testis |
| 9823 | SRS1198656 | liver |
| 9823 | SRS1198657 | liver |
| 9823 | SRS1198658 | liver |
| 9823 | SRS1792915 | brain |
| 9823 | SRS1930476 | heart |

| | | |
|---|---|---|
| 9823 | SRS1930478 | heart |
| 9823 | SRS1930481 | heart |
| 9823 | SRS1930492 | heart |
| 9823 | SRS1930495 | heart |
| 9823 | SRS1930531 | kidney |
| 9823 | SRS1930532 | kidney |
| 9823 | SRS1930533 | kidney |
| 9823 | SRS1930588 | kidney |
| 9823 | SRS1930589 | kidney |
| 9823 | SRS1930592 | kidney |
| 9823 | SRS2140178 | brain |
| 9823 | SRS2140180 | brain |
| 9823 | SRS2140181 | brain |
| 9823 | SRS2140182 | brain |
| 9823 | SRS2140183 | brain |
| 9823 | SRS2140184 | brain |
| 9823 | SRS2518416 | liver |
| 9823 | SRS2518419 | liver |
| 9823 | SRS2518420 | liver |
| 9823 | SRS2518423 | liver |
| 9823 | SRS2602179 | ovary |
| 9823 | SRS2602181 | ovary |
| 9823 | SRS2602183 | ovary |
| 9823 | SRS2659243 | liver |
| 9823 | SRS2659244 | liver |
| 9823 | SRS2659245 | liver |
| 9823 | SRS2659246 | liver |
| 9823 | SRS2659248 | liver |
| 9823 | SRS2760027 | ovary |
| 9823 | SRS2760028 | ovary |

| | | |
|---|---|---|
| 9823 | SRS2760029 | ovary |
| 9823 | SRS2760030 | ovary |
| 9823 | SRS2760031 | ovary |
| 9823 | SRS2760032 | ovary |
| 9823 | SRS380782 | kidney |
| 9823 | SRS380783 | kidney |
| 9823 | SRS380827 | brain |
| 9823 | SRS380828 | brain |
| 9823 | SRS388205 | heart |
| 9823 | SRS388208 | kidney |
| 9823 | SRS529859 | liver |
| 9823 | SRS529863 | liver |
| 9823 | SRS529865 | liver |
| 9796 | SRR12641943 | Liver |
| 9796 | SRR12641945 | Heart |
| 9796 | SRR12641949 | Liver |
| 9796 | SRR12641952 | Heart |
| 9796 | SRR12641955 | Liver |
| 9796 | SRR12641957 | Heart |
| 9796 | SRR12641960 | Liver |
| 9796 | SRR12641961 | Heart |
| 9796 | SRR12641962 | Liver |
| 9796 | SRR12641965 | Liver |
| 9796 | SRR12641967 | Heart |
| 9796 | SRR12641970 | Liver |
| 9796 | SRR12641972 | Heart |
| 9796 | SRR12641974 | Heart |
| 9796 | SRR21046748 | Liver |
| 9796 | SRR21046749 | Liver |
| 9796 | SRR21046750 | Liver |

| | | |
|---|---|---|
| 9796 | SRR21046751 | Liver |
| 9796 | SRR21046752 | Liver |
| 9796 | SRR21046753 | Liver |
| 9796 | SRR21046754 | Liver |
| 9796 | SRR21046755 | Liver |
| 9796 | SRR3403817 | Brain |
| 9796 | SRR3403818 | Brain |
| 9796 | SRR3403819 | Brain |
| 9796 | SRR3403821 | Brain |
| 9796 | SRR3403822 | Brain |
| 9796 | SRR3403825 | Brain |
| 9796 | SRR3403826 | Brain |
| 9796 | SRR3403827 | Brain |
| 9796 | SRR3403828 | Brain |
| 9796 | SRR6361828 | Testis |
| 9796 | SRR6361829 | Testis |
| 9796 | SRR636899 | Kidney |
| 9796 | SRR636900 | Kidney |
| 9796 | SRR636901 | Kidney |
| 9796 | SRR8303651 | Testis |
| 9796 | SRR8303652 | Testis |
| 9796 | SRR8303653 | Testis |
| 9796 | SRR8303654 | Testis |
| 9796 | SRR8303655 | Testis |
| 9796 | SRR8303656 | Testis |
| 9796 | SRR8303657 | Testis |
| 9796 | SRR8303658 | Testis |
| 9796 | SRR8303659 | Testis |
| 9796 | SRR8303660 | Testis |
| 9685 | SRR11852802 | Ovary |

| | | |
|---|---|---|
| 9685 | SRR11852803 | Ovary |
| 9685 | SRR11852804 | Ovary |
| 9685 | SRR11852805 | Ovary |
| 9685 | SRR11852806 | Ovary |
| 9685 | SRR11852807 | Ovary |
| 9685 | SRR11852808 | Ovary |
| 9685 | SRR11852809 | Ovary |
| 9685 | SRR11852810 | Ovary |
| 9685 | SRR11939407 | Heart |
| 9685 | SRR11939411 | Liver |
| 9685 | SRR11939412 | Kidney |
| 9685 | SRR11939414 | Heart |
| 9685 | SRR11939419 | Liver |
| 9685 | SRR11939420 | Kidney |
| 9685 | SRR11939422 | Heart |
| 9685 | SRR11939457 | Liver |
| 9685 | SRR11939468 | Kidney |
| 9685 | SRR12613016 | Ovary |
| 9685 | SRR12613017 | Ovary |
| 9685 | SRR12613018 | Ovary |
| 9685 | SRR12613023 | Ovary |
| 9685 | SRR12613034 | Ovary |
| 9685 | SRR12613035 | Ovary |
| 9685 | SRR15011234 | Testis |
| 9685 | SRR15011235 | Testis |
| 9685 | SRR15011236 | Testis |
| 9685 | SRR15011242 | Testis |
| 9685 | SRR15011243 | Testis |
| 9685 | SRR15011244 | Testis |
| 9685 | SRR15011245 | Testis |

| | | |
|---|---|---|
| 9685 | SRR15011246 | Testis |
| 9685 | SRR15011247 | Testis |
| 9685 | SRR15011248 | Testis |
| 9685 | SRR21657489 | Brain |
| 9685 | SRR21657490 | Brain |
| 9685 | SRR21657498 | Brain |
| 9685 | SRR21657509 | Brain |
| 9685 | SRR21657516 | Liver |
| 9685 | SRR21657517 | Liver |
| 9685 | SRR21657518 | Liver |
| 9685 | SRR21657519 | Liver |
| 9685 | SRR21657521 | Liver |
| 9685 | SRR21657522 | Liver |
| 9685 | SRR21657523 | Kidney |
| 9685 | SRR21657524 | Kidney |
| 9685 | SRR21657525 | Kidney |
| 9685 | SRR21657526 | Kidney |
| 9685 | SRR21657527 | Kidney |
| 9685 | SRR21657528 | Kidney |
| 9685 | SRR21657529 | Kidney |
| 9685 | SRR21657530 | Kidney |
| 9685 | SRR21657532 | Kidney |
| 9685 | SRR21657533 | Kidney |
| 9685 | SRR21657534 | Kidney |
| 9685 | SRR21657535 | Kidney |
| 9685 | SRR21657549 | Heart |
| 9685 | SRR21657550 | Heart |
| 9685 | SRR21657551 | Heart |
| 9685 | SRR21657552 | Heart |
| 9685 | SRR21657554 | Heart |

| | | |
|---|---|---|
| 9685 | SRR21657555 | Heart |
| 9685 | SRR21657562 | Brain |
| 9685 | SRR21657563 | Brain |
| 9646 | SRR10215705 | Heart |
| 9646 | SRR10215706 | Heart |
| 9646 | SRR10215707 | Heart |
| 9646 | SRR10215708 | Heart |
| 9646 | SRR10215709 | Heart |
| 9646 | SRR10215710 | Liver |
| 9646 | SRR10215711 | Liver |
| 9646 | SRR10215712 | Liver |
| 9646 | SRR10215713 | Liver |
| 9646 | SRR10215714 | Liver |
| 9646 | SRR10215725 | Kidney |
| 9646 | SRR10215726 | Kidney |
| 9646 | SRR10215727 | Kidney |
| 9646 | SRR10215728 | Kidney |
| 9646 | SRR10215729 | Kidney |
| 9646 | SRR15569469 | Testis |
| 9646 | SRR15569470 | Testis |
| 9646 | SRR15569471 | Testis |
| 9646 | SRR15569473 | Testis |
| 9646 | SRR15569474 | Testis |
| 9646 | SRR15569475 | Testis |
| 9646 | SRR15569482 | Testis |
| 9646 | SRR15569483 | Testis |
| 9646 | SRR15569486 | Testis |
| 9646 | SRR15569493 | Testis |
| 9646 | SRR15569494 | Testis |
| 9646 | SRR15569495 | Testis |

| | | |
|---|---|---|
| 9646 | SRR15569496 | Testis |
| 9646 | SRR15569497 | Testis |
| 9646 | SRR15569498 | Testis |
| 9646 | SRR2307714 | Brain |
| 9646 | SRR2307789 | Brain |
| 9646 | SRR2307886 | Ovary |
| 9646 | SRR2308103 | Liver |
| 9646 | SRR7611054 | Brain |
| 9646 | SRR7611055 | Liver |
| 9615 | ERS1090466 | liver |
| 9615 | ERS1090467 | liver |
| 9615 | ERS1090468 | liver |
| 9615 | ERS1446917 | testis |
| 9615 | ERS1446918 | testis |
| 9615 | ERS1446919 | testis |
| 9615 | ERS1446920 | testis |
| 9615 | ERS1446922 | testis |
| 9615 | ERS1446923 | testis |
| 9615 | ERS1446924 | testis |
| 9615 | ERS1446925 | testis |
| 9615 | SRR11939292 | Kidney |
| 9615 | SRR11939348 | Kidney |
| 9615 | SRR11939371 | Kidney |
| 9615 | SRR1207321 | Testis |
| 9615 | SRR12544780 | Liver |
| 9615 | SRR12544781 | Liver |
| 9615 | SRR12544782 | Liver |
| 9615 | SRR12544783 | Liver |
| 9615 | SRR12544784 | Liver |
| 9615 | SRR12544785 | Liver |

| | | |
|---|---|---|
| 9615 | SRR12544786 | Liver |
| 9615 | SRR12544787 | Liver |
| 9615 | SRR12544788 | Liver |
| 9615 | SRR12544789 | Liver |
| 9615 | SRR12544790 | Liver |
| 9615 | SRR12544791 | Liver |
| 9615 | SRR12544792 | Liver |
| 9615 | SRR12544793 | Liver |
| 9615 | SRR12544794 | Liver |
| 9615 | SRR16767722 | Testis |
| 9615 | SRR16767723 | Testis |
| 9615 | SRR16767724 | Testis |
| 9615 | SRR16767725 | Testis |
| 9615 | SRR16767728 | Testis |
| 9615 | SRR16767729 | Testis |
| 9615 | SRR18300827 | Brain |
| 9615 | SRR18300828 | Brain |
| 9615 | SRR18300829 | Brain |
| 9615 | SRR18300830 | Brain |
| 9615 | SRR18300831 | Brain |
| 9615 | SRR18300832 | Brain |
| 9615 | SRR18300833 | Brain |
| 9615 | SRR18300834 | Brain |
| 9615 | SRR18300835 | Brain |
| 9615 | SRR18300836 | Brain |
| 9615 | SRR18300837 | Brain |
| 9615 | SRR18300838 | Brain |
| 9615 | SRR18300839 | Brain |
| 9615 | SRR5889316 | Heart |
| 9615 | SRR5889319 | Heart |

| | | |
|---|---|---|
| 9615 | SRR5889325 | Heart |
| 9615 | SRR5889332 | Heart |
| 9615 | SRR5889335 | Heart |
| 9615 | SRR5889336 | Heart |
| 9615 | SRR5889342 | Heart |
| 9615 | SRR5889343 | Heart |
| 9615 | SRR6206926 | Testis |
| 9615 | SRR8996963 | Heart |
| 9615 | SRR8996964 | Heart |
| 9615 | SRR8996969 | Heart |
| 9615 | SRR8996973 | Heart |
| 9615 | SRR8996974 | Heart |
| 9615 | SRR8996975 | Heart |
| 9615 | SRR8996978 | Heart |
| 9615 | SRS072646 | brain |
| 9615 | SRS072647 | liver |
| 9615 | SRS072744 | heart |
| 9615 | SRS072745 | ovary |
| 9615 | SRS072747 | testis |
| 9615 | SRS072786 | kidney |
| 9615 | SRS1995919 | heart |
| 9615 | SRS1995925 | heart |
| 9615 | SRS1995929 | heart |
| 9615 | SRS1995935 | heart |
| 9615 | SRS1995939 | heart |
| 9615 | SRS1995940 | heart |
| 9615 | SRS1995955 | heart |
| 9615 | SRS1995959 | heart |
| 9615 | SRS1995960 | heart |
| 9615 | SRS1995961 | heart |

| | | |
|---|---|---|
| 9615 | SRS1995965 | heart |
| 9615 | SRS2040965 | kidney |
| 9615 | SRS2040966 | kidney |
| 9615 | SRS2401093 | liver |
| 9615 | SRS2401097 | heart |
| 9615 | SRS2401100 | liver |
| 9615 | SRS2401103 | heart |
| 9615 | SRS2401106 | kidney |
| 9615 | SRS2401108 | liver |
| 9615 | SRS2401109 | kidney |
| 9615 | SRS2401110 | heart |
| 9615 | SRS2401112 | liver |
| 9615 | SRS2401115 | liver |
| 9615 | SRS332229 | brain |
| 9615 | SRS332231 | kidney |
| 9615 | SRS380696 | liver |
| 9615 | SRS380697 | liver |
| 9615 | SRS380746 | kidney |
| 9615 | SRS380747 | kidney |
| 9615 | SRS380791 | brain |
| 9615 | SRS380792 | brain |
| 9601 | ERR3491602 | Brain |
| 9601 | ERR3491608 | Brain |
| 9601 | ERR3491609 | Brain |
| 9601 | ERR3491610 | Brain |
| 9601 | ERR3491615 | Brain |
| 9601 | ERR3491616 | Brain |
| 9601 | ERR3491621 | Brain |
| 9601 | ERR3491623 | Brain |
| 9601 | ERR3491624 | Brain |

| | | |
|---|---|---|
| 9601 | ERR3491627 | Brain |
| 9601 | ERR3491628 | Brain |
| 9601 | ERR3491630 | Brain |
| 9601 | ERR3491634 | Brain |
| 9601 | ERR3491636 | Brain |
| 9601 | ERR3491637 | Brain |
| 9598 | SRR2040586 | Heart |
| 9598 | SRR2040587 | Heart |
| 9598 | SRR2040588 | Liver |
| 9598 | SRR2040589 | Liver |
| 9598 | SRR2040590 | Testis |
| 9598 | SRR2040591 | Testis |
| 9598 | SRR8750641 | Brain |
| 9598 | SRR8750651 | Brain |
| 9598 | SRR8750671 | Brain |
| 9598 | SRR8750672 | Brain |
| 9598 | SRR8750674 | Brain |
| 9598 | SRR8750680 | Brain |
| 9598 | SRR8750681 | Brain |
| 9598 | SRR8750684 | Brain |
| 9598 | SRR8750693 | Brain |
| 9598 | SRR8750704 | Brain |
| 9598 | SRR8750712 | Brain |
| 9598 | SRR8750714 | Brain |
| 9598 | SRR8750716 | Brain |
| 9598 | SRR8750717 | Brain |
| 9598 | SRR8750718 | Brain |
| 9597 | SRR8750398 | Brain |
| 9597 | SRR8750416 | Brain |
| 9597 | SRR8750420 | Brain |

| | | |
|---|---|---|
| 9597 | SRR8750426 | Brain |
| 9597 | SRR8750430 | Brain |
| 9597 | SRR8750436 | Brain |
| 9597 | SRR8750440 | Brain |
| 9597 | SRR8750445 | Brain |
| 9597 | SRR8750604 | Brain |
| 9597 | SRR8750606 | Brain |
| 9597 | SRR8750608 | Brain |
| 9597 | SRR8750616 | Brain |
| 9597 | SRR8750621 | Brain |
| 9597 | SRR8750625 | Brain |
| 9597 | SRR8750628 | Brain |
| 9595 | SRR5804494 | Brain |
| 9595 | SRR5804495 | Brain |
| 9595 | SRR5804496 | Brain |
| 9595 | SRR5804497 | Brain |
| 9595 | SRR5804498 | Brain |
| 9595 | SRR5804499 | Brain |
| 9595 | SRR5804500 | Brain |
| 9595 | SRR5804501 | Brain |
| 9595 | SRR5804502 | Brain |
| 9595 | SRR5804503 | Brain |
| 9595 | SRR5804504 | Brain |
| 9595 | SRR5804506 | Brain |
| 9595 | SRR5804507 | Brain |
| 9595 | SRR5804508 | Brain |
| 9595 | SRR5804509 | Brain |
| 9555 | SRR1041110 | Brain |
| 9555 | SRR1041111 | Brain |
| 9555 | SRR1041113 | Heart |

| | | |
|---|---|---|
| 9555 | SRR1041114 | Kidney |
| 9555 | SRR1041115 | Liver |
| 9555 | SRR1045084 | Brain |
| 9555 | SRR1045085 | Brain |
| 9555 | SRR1045089 | Brain |
| 9544 | ERS1090480 | liver |
| 9544 | ERS1090481 | liver |
| 9544 | ERS1090482 | liver |
| 9544 | ERS1090483 | liver |
| 9544 | SRR10765326 | Testis |
| 9544 | SRR10765327 | Testis |
| 9544 | SRR10765328 | Testis |
| 9544 | SRR10765329 | Testis |
| 9544 | SRR10765330 | Testis |
| 9544 | SRR10765331 | Testis |
| 9544 | SRR10765332 | Testis |
| 9544 | SRR10765333 | Testis |
| 9544 | SRR10765334 | Testis |
| 9544 | SRR10765335 | Testis |
| 9544 | SRR11939363 | Heart |
| 9544 | SRR11939368 | Kidney |
| 9544 | SRR11939369 | Heart |
| 9544 | SRR11939375 | Kidney |
| 9544 | SRR11939377 | Heart |
| 9544 | SRR11939383 | Kidney |
| 9544 | SRR13083225 | Kidney |
| 9544 | SRR13083226 | Kidney |
| 9544 | SRR13083227 | Kidney |
| 9544 | SRR2040593 | Heart |
| 9544 | SRR223519 | Heart |

| | | |
|---|---|---|
| 9544 | SRR5448350 | Testis |
| 9544 | SRR5448351 | Testis |
| 9544 | SRR5448353 | Testis |
| 9544 | SRR5448354 | Testis |
| 9544 | SRR5448355 | Testis |
| 9544 | SRR594457 | Heart |
| 9544 | SRR6056757 | Heart |
| 9544 | SRR6056758 | Kidney |
| 9544 | SRR6056763 | Heart |
| 9544 | SRR6056764 | Kidney |
| 9544 | SRR6074391 | Liver |
| 9544 | SRR6074419 | Liver |
| 9544 | SRR6074420 | Liver |
| 9544 | SRR6074421 | Liver |
| 9544 | SRR6074422 | Liver |
| 9544 | SRR6074423 | Liver |
| 9544 | SRR6074424 | Liver |
| 9544 | SRR6074425 | Liver |
| 9544 | SRR6074426 | Liver |
| 9544 | SRR6074427 | Liver |
| 9544 | SRR6074448 | Liver |
| 9544 | SRR6074449 | Liver |
| 9544 | SRR6074450 | Liver |
| 9544 | SRR6074451 | Liver |
| 9544 | SRR6074452 | Liver |
| 9544 | SRR8750509 | Brain |
| 9544 | SRR8750513 | Brain |
| 9544 | SRR8750521 | Brain |
| 9544 | SRR8750530 | Brain |
| 9544 | SRR8750540 | Brain |

| | | |
|---|---|---|
| 9544 | SRR8750544 | Brain |
| 9544 | SRR8750545 | Brain |
| 9544 | SRR8750549 | Brain |
| 9544 | SRR8750567 | Brain |
| 9544 | SRR8750572 | Brain |
| 9544 | SRR8750589 | Brain |
| 9544 | SRR8750590 | Brain |
| 9544 | SRR8750591 | Brain |
| 9544 | SRR8750592 | Brain |
| 9544 | SRR8750593 | Brain |
| 9544 | SRS1992612 | heart |
| 9544 | SRS1992614 | kidney |
| 9544 | SRS1992618 | liver |
| 9544 | SRS211172 | brain |
| 9544 | SRS211173 | heart |
| 9544 | SRS211174 | kidney |
| 9544 | SRS211175 | liver |
| 9544 | SRS2124532 | testis |
| 9544 | SRS2124533 | testis |
| 9544 | SRS2124534 | testis |
| 9544 | SRS2124535 | testis |
| 9544 | SRS2124536 | testis |
| 9544 | SRS2124537 | testis |
| 9544 | SRS2124538 | testis |
| 9544 | SRS2124539 | testis |
| 9544 | SRS2124540 | testis |
| 9544 | SRS2124541 | testis |
| 9544 | SRS282392 | liver |
| 9544 | SRS282394 | testis |
| 9544 | SRS369754 | brain |

| | | |
|---|---|---|
| 9544 | SRS369756 | heart |
| 9544 | SRS369757 | kidney |
| 9544 | SRS369758 | liver |
| 9544 | SRS369762 | testis |
| 9544 | SRS369763 | brain |
| 9544 | SRS369765 | heart |
| 9544 | SRS369766 | kidney |
| 9544 | SRS369767 | liver |
| 9544 | SRS369771 | testis |
| 9544 | SRS369772 | brain |
| 9544 | SRS369774 | heart |
| 9544 | SRS369775 | kidney |
| 9544 | SRS369776 | liver |
| 9544 | SRS369780 | testis |
| 9544 | SRS379370 | kidney |
| 9544 | SRS576978 | heart |
| 9544 | SRS716802 | heart |
| 9544 | SRS716803 | kidney |
| 9544 | SRS716804 | liver |
| 9544 | SRS716810 | ovary |
| 9544 | SRS945659 | testis |
| 9544 | SRS945660 | liver |
| 9544 | SRS945661 | heart |
| 9544 | SRS945662 | brain |
| 9541 | SRR14825551 | Liver |
| 9541 | SRR14825554 | Brain |
| 9541 | SRR14825557 | Liver |
| 9541 | SRR14825558 | Brain |
| 9541 | SRR14825559 | Brain |
| 9541 | SRR14825560 | Brain |

| | | |
|---|---|---|
| 9541 | SRR14825572 | Liver |
| 9541 | SRR14825573 | Brain |
| 9541 | SRR14825574 | Brain |
| 9541 | SRR14825575 | Brain |
| 9541 | SRR14825589 | Liver |
| 9541 | SRR14825590 | Brain |
| 9541 | SRR14825591 | Brain |
| 9541 | SRR14825592 | Brain |
| 9541 | SRR14825594 | Liver |
| 9541 | SRR14825595 | Brain |
| 9541 | SRR14825596 | Brain |
| 9541 | SRR14825597 | Brain |
| 9541 | SRR14825600 | Liver |
| 9541 | SRR14825601 | Brain |
| 9541 | SRR14825603 | Brain |
| 9541 | SRR15905854 | Testis |
| 9541 | SRR15905867 | Testis |
| 9541 | SRR15905868 | Testis |
| 9541 | SRR15905869 | Testis |
| 9541 | SRR15905884 | Testis |
| 9541 | SRR15905885 | Testis |
| 9541 | SRR15905911 | Testis |
| 9541 | SRR18674643 | Liver |
| 9541 | SRR18674654 | Liver |
| 9541 | SRR18674655 | Liver |
| 9541 | SRR18674656 | Liver |
| 9541 | SRR18674657 | Liver |
| 9541 | SRR18674658 | Liver |
| 9541 | SRR18674659 | Liver |
| 9541 | SRR18674661 | Liver |

| | | |
|---|---|---|
| 9541 | SRR18674662 | Liver |
| 9541 | SRR8474208 | Testis |
| 9541 | SRR8474209 | Testis |
| 9541 | SRR8474210 | Testis |
| 9541 | SRR8474211 | Testis |
| 9541 | SRR8474212 | Testis |
| 9541 | SRR8474213 | Testis |
| 9541 | SRR8474214 | Testis |
| 9541 | SRR8474215 | Testis |
| 9483 | ERS1090469 | liver |
| 9483 | ERS1090471 | liver |
| 9483 | ERS1090516 | liver |
| 9483 | ERS1090517 | liver |
| 9483 | SRR10466922 | Brain |
| 9483 | SRR10466924 | Brain |
| 9483 | SRR10466927 | Brain |
| 9483 | SRR10466928 | Brain |
| 9483 | SRR10466929 | Brain |
| 9483 | SRR1758977 | Brain |
| 9483 | SRR1758979 | Brain |
| 9483 | SRR1758981 | Heart |
| 9483 | SRR1758982 | Heart |
| 9483 | SRR1758983 | Kidney |
| 9483 | SRR1758984 | Liver |
| 9483 | SRR19268427 | Brain |
| 9483 | SRR19268428 | Brain |
| 9483 | SRR19268431 | Brain |
| 9483 | SRR19268432 | Brain |
| 9483 | SRR19268436 | Brain |
| 9483 | SRR19268437 | Brain |

| | | |
|---|---|---|
| 9483 | SRR19268438 | Brain |
| 9483 | SRR19268439 | Brain |
| 9483 | SRR5928355 | Heart |
| 9483 | SRR5928360 | Liver |
| 9483 | SRR5928361 | Kidney |
| 9483 | SRS2426394 | kidney |
| 9483 | SRS2426397 | brain |
| 9483 | SRS2426398 | liver |
| 9483 | SRS2426399 | heart |
| 9483 | SRS470452 | testis |
| 9483 | SRS470454 | liver |
| 9483 | SRS470552 | kidney |
| 9483 | SRS470553 | heart |
| 9483 | SRS470554 | brain |
| 9483 | SRS819378 | brain |
| 9483 | SRS819379 | brain |
| 9483 | SRS819381 | heart |
| 9483 | SRS819382 | kidney |
| 9483 | SRS819383 | liver |
| 9103 | SRR10746605 | Testis |
| 9103 | SRR10746606 | Testis |
| 9103 | SRR10746607 | Testis |
| 9103 | SRR10746608 | Testis |
| 9103 | SRR10746609 | Testis |
| 9103 | SRR10746611 | Testis |
| 9103 | SRR10746612 | Testis |
| 9103 | SRR10746613 | Testis |
| 9103 | SRR10746614 | Testis |
| 9103 | SRR10746615 | Testis |
| 9103 | SRR10746616 | Testis |

| | | |
|---|---|---|
| 9103 | SRR10746617 | Testis |
| 9103 | SRR10746618 | Testis |
| 9103 | SRR10746619 | Testis |
| 9103 | SRR4244331 | Liver |
| 9103 | SRR4244334 | Liver |
| 9103 | SRR4244337 | Liver |
| 9103 | SRR4244341 | Liver |
| 9103 | SRR4244357 | Liver |
| 9103 | SRR4244361 | Liver |
| 9103 | SRR4244364 | Liver |
| 9103 | SRR4244367 | Liver |
| 9103 | SRR9044387 | Brain |
| 9103 | SRR9044404 | Brain |
| 9103 | SRR9044415 | Brain |
| 9103 | SRR9044428 | Brain |
| 9103 | SRR9044430 | Brain |
| 9103 | SRR9044431 | Brain |
| 9103 | SRR9044433 | Brain |
| 9103 | SRR9044434 | Brain |
| 9103 | SRR9044435 | Brain |
| 9103 | SRR9044436 | Brain |
| 9103 | SRR9044437 | Brain |
| 9103 | SRR9044438 | Brain |
| 9103 | SRR9044439 | Brain |
| 9103 | SRR9044440 | Brain |
| 9103 | SRR9044441 | Brain |
| 9031 | ERS1070003 | ovary |
| 9031 | ERS2006582 | liver |
| 9031 | ERS2006585 | liver |
| 9031 | ERS2006593 | liver |

| | | |
|---|---|---|
| 9031 | ERS2006594 | liver |
| 9031 | ERS353481 | testis |
| 9031 | ERS353485 | ovary |
| 9031 | ERS353500 | heart |
| 9031 | ERS353503 | heart |
| 9031 | SRR10870449 | Testis |
| 9031 | SRR10870450 | Testis |
| 9031 | SRR10870451 | Testis |
| 9031 | SRR10870452 | Testis |
| 9031 | SRR10870453 | Testis |
| 9031 | SRR10870454 | Testis |
| 9031 | SRR10870455 | Testis |
| 9031 | SRR10870456 | Testis |
| 9031 | SRR11782411 | Heart |
| 9031 | SRR11782412 | Heart |
| 9031 | SRR11782413 | Heart |
| 9031 | SRR11782414 | Heart |
| 9031 | SRR11782415 | Heart |
| 9031 | SRR11782416 | Heart |
| 9031 | SRR11782417 | Heart |
| 9031 | SRR11782418 | Heart |
| 9031 | SRR11908628 | Heart |
| 9031 | SRR11908629 | Heart |
| 9031 | SRR11908630 | Heart |
| 9031 | SRR11928976 | Heart |
| 9031 | SRR11928977 | Heart |
| 9031 | SRR11928978 | Heart |
| 9031 | SRR11928979 | Heart |
| 9031 | SRR11939389 | Kidney |
| 9031 | SRR11939397 | Kidney |

| | | |
|---|---|---|
| 9031 | SRR11939405 | Kidney |
| 9031 | SRR14325372 | Brain |
| 9031 | SRR14325373 | Ovary |
| 9031 | SRR14325374 | Brain |
| 9031 | SRR14325375 | Ovary |
| 9031 | SRR14325376 | Brain |
| 9031 | SRR14325377 | Ovary |
| 9031 | SRR14325378 | Brain |
| 9031 | SRR14325379 | Ovary |
| 9031 | SRR14325380 | Brain |
| 9031 | SRR14325381 | Ovary |
| 9031 | SRR14325382 | Brain |
| 9031 | SRR14325383 | Brain |
| 9031 | SRR14325384 | Ovary |
| 9031 | SRR14325385 | Brain |
| 9031 | SRR14325386 | Ovary |
| 9031 | SRR14325387 | Brain |
| 9031 | SRR14325388 | Ovary |
| 9031 | SRR14325389 | Brain |
| 9031 | SRR14325390 | Ovary |
| 9031 | SRR14325391 | Brain |
| 9031 | SRR14325392 | Ovary |
| 9031 | SRR14325393 | Ovary |
| 9031 | SRR14325394 | Brain |
| 9031 | SRR14325395 | Ovary |
| 9031 | SRR14325396 | Brain |
| 9031 | SRR14325397 | Ovary |
| 9031 | SRR14325398 | Brain |
| 9031 | SRR14325399 | Ovary |
| 9031 | SRR14325400 | Brain |

| | | |
|---|---|---|
| 9031 | SRR14325401 | Ovary |
| 9031 | SRR19880159 | Testis |
| 9031 | SRR19880160 | Testis |
| 9031 | SRR19880161 | Testis |
| 9031 | SRR19880162 | Testis |
| 9031 | SRR19880163 | Testis |
| 9031 | SRR19880164 | Testis |
| 9031 | SRR20821354 | Liver |
| 9031 | SRR20821355 | Liver |
| 9031 | SRR20821358 | Liver |
| 9031 | SRR20821360 | Liver |
| 9031 | SRR20821362 | Liver |
| 9031 | SRR20821363 | Liver |
| 9031 | SRR20821368 | Liver |
| 9031 | SRR20821375 | Liver |
| 9031 | SRR20821376 | Liver |
| 9031 | SRR20821381 | Liver |
| 9031 | SRR20821382 | Liver |
| 9031 | SRR20821384 | Liver |
| 9031 | SRR20821386 | Liver |
| 9031 | SRR20821387 | Liver |
| 9031 | SRR20821388 | Liver |
| 9031 | SRR4933176 | Kidney |
| 9031 | SRR4933177 | Kidney |
| 9031 | SRR4933178 | Kidney |
| 9031 | SRR4933179 | Kidney |
| 9031 | SRR4933180 | Kidney |
| 9031 | SRR4933181 | Kidney |
| 9031 | SRR4933182 | Kidney |
| 9031 | SRR9606516 | Kidney |

| | | |
|---|---|---|
| 9031 | SRR9606524 | Kidney |
| 9031 | SRR9606532 | Kidney |
| 9031 | SRR9606540 | Kidney |
| 9031 | SRR9606548 | Kidney |
| 9031 | SRS1114756 | testis |
| 9031 | SRS1114757 | testis |
| 9031 | SRS1783075 | kidney |
| 9031 | SRS1783076 | kidney |
| 9031 | SRS1783077 | kidney |
| 9031 | SRS1783078 | kidney |
| 9031 | SRS1783079 | kidney |
| 9031 | SRS1930484 | heart |
| 9031 | SRS1930487 | heart |
| 9031 | SRS1930490 | heart |
| 9031 | SRS1930517 | heart |
| 9031 | SRS1930520 | heart |
| 9031 | SRS1930523 | heart |
| 9031 | SRS1930585 | kidney |
| 9031 | SRS1930586 | kidney |
| 9031 | SRS1930587 | kidney |
| 9031 | SRS1930594 | kidney |
| 9031 | SRS1930595 | kidney |
| 9031 | SRS1930596 | kidney |
| 9031 | SRS2136320 | liver |
| 9031 | SRS2136321 | liver |
| 9031 | SRS2136322 | liver |
| 9031 | SRS2136323 | liver |
| 9031 | SRS2136324 | liver |
| 9031 | SRS2986822 | liver |
| 9031 | SRS3052303 | liver |

| | | |
|---|---|---|
| 9031 | SRS3052304 | liver |
| 9031 | SRS3074381 | heart |
| 9031 | SRS359046 | ovary |
| 9031 | SRS366864 | brain |
| 9031 | SRS366865 | liver |
| 9031 | SRS366866 | kidney |
| 9031 | SRS366867 | heart |
| 9031 | SRS369808 | brain |
| 9031 | SRS369810 | heart |
| 9031 | SRS369811 | kidney |
| 9031 | SRS369812 | liver |
| 9031 | SRS369816 | testis |
| 9031 | SRS369817 | brain |
| 9031 | SRS369819 | heart |
| 9031 | SRS369820 | kidney |
| 9031 | SRS369821 | liver |
| 9031 | SRS369825 | testis |
| 9031 | SRS369828 | heart |
| 9031 | SRS369829 | kidney |
| 9031 | SRS369834 | testis |
| 8479 | SRR13381391 | Liver |
| 8479 | SRR13381392 | Liver |
| 8479 | SRR13381393 | Liver |
| 8479 | SRR13381394 | Liver |
| 8479 | SRR13381397 | Liver |
| 8479 | SRR13381398 | Liver |
| 8479 | SRR8718961 | Heart |
| 8479 | SRR8718962 | Heart |
| 8479 | SRR8718963 | Heart |
| 8479 | SRR8718964 | Heart |

| | | |
|---|---|---|
| 8479 | SRR8718965 | Heart |
| 8479 | SRR8718966 | Heart |
| 8479 | SRR8718967 | Heart |
| 8479 | SRR8718968 | Heart |
| 8479 | SRR8718969 | Heart |
| 8479 | SRR8718970 | Heart |
| 8479 | SRR8718971 | Heart |
| 8479 | SRR8718972 | Heart |
| 8479 | SRR8718973 | Heart |
| 8479 | SRR8718974 | Heart |
| 8479 | SRR8718975 | Heart |
| 8364 | SRR12368258 | Brain |
| 8364 | SRR12368259 | Brain |
| 8364 | SRR12368260 | Brain |
| 8364 | SRR12368261 | Brain |
| 8364 | SRR12368262 | Brain |
| 8364 | SRR12368263 | Brain |
| 8364 | SRR12368264 | Brain |
| 8364 | SRR12368265 | Brain |
| 8364 | SRR12368266 | Brain |
| 8364 | SRR12368267 | Brain |
| 8364 | SRR12368268 | Brain |
| 8364 | SRR12368269 | Brain |
| 8364 | SRR12368270 | Brain |
| 8364 | SRR12368271 | Brain |
| 8364 | SRR12368272 | Brain |
| 8364 | SRS2073206 | liver |
| 8364 | SRS366873 | brain |
| 8364 | SRS366874 | liver |
| 8364 | SRS366875 | kidney |

| | | |
|---|---|---|
| 8364 | SRS366876 | heart |
| 8364 | SRS568433 | brain |
| 8364 | SRS568436 | kidney |
| 7955 | ERS017858 | kidney |
| 7955 | ERS017860 | heart |
| 7955 | ERS017861 | brain |
| 7955 | SRR10272861 | Testis |
| 7955 | SRR10854654 | Testis |
| 7955 | SRR10854656 | Testis |
| 7955 | SRR11033311 | Heart |
| 7955 | SRR11033312 | Heart |
| 7955 | SRR11033313 | Heart |
| 7955 | SRR11869424 | Heart |
| 7955 | SRR11869425 | Heart |
| 7955 | SRR11869426 | Heart |
| 7955 | SRR12446287 | Brain |
| 7955 | SRR12446289 | Brain |
| 7955 | SRR12446290 | Brain |
| 7955 | SRR12446292 | Brain |
| 7955 | SRR12446294 | Brain |
| 7955 | SRR12446295 | Brain |
| 7955 | SRR14693495 | Liver |
| 7955 | SRR14693498 | Liver |
| 7955 | SRR14693499 | Liver |
| 7955 | SRR15712664 | Kidney |
| 7955 | SRR15712671 | Kidney |
| 7955 | SRR15712674 | Kidney |
| 7955 | SRR17026704 | Brain |
| 7955 | SRR17026705 | Brain |
| 7955 | SRR17026706 | Brain |

| | | |
|---|---|---|
| 7955 | SRR17026707 | Brain |
| 7955 | SRR17026708 | Brain |
| 7955 | SRR17026709 | Brain |
| 7955 | SRR17026710 | Brain |
| 7955 | SRR17026711 | Brain |
| 7955 | SRR17026712 | Brain |
| 7955 | SRR17248539 | Kidney |
| 7955 | SRR17248540 | Kidney |
| 7955 | SRR17248541 | Kidney |
| 7955 | SRR17259773 | Kidney |
| 7955 | SRR17259774 | Kidney |
| 7955 | SRR17259775 | Kidney |
| 7955 | SRR17259776 | Kidney |
| 7955 | SRR17259777 | Kidney |
| 7955 | SRR17259778 | Kidney |
| 7955 | SRR18297397 | Liver |
| 7955 | SRR18297399 | Liver |
| 7955 | SRR18297400 | Liver |
| 7955 | SRR19387525 | Kidney |
| 7955 | SRR19387526 | Kidney |
| 7955 | SRR19387527 | Kidney |
| 7955 | SRR19831100 | Liver |
| 7955 | SRR19831102 | Liver |
| 7955 | SRR20018673 | Liver |
| 7955 | SRR20018674 | Liver |
| 7955 | SRR20018675 | Liver |
| 7955 | SRR5378555 | Testis |
| 7955 | SRR5378556 | Testis |
| 7955 | SRR5378557 | Testis |
| 7955 | SRR5378566 | Testis |

| | | |
|---|---|---|
| 7955 | SRR5378567 | Testis |
| 7955 | SRR5378568 | Testis |
| 7955 | SRR6257756 | Testis |
| 7955 | SRR6257759 | Testis |
| 7955 | SRR6257760 | Testis |
| 7955 | SRR6257761 | Testis |
| 7955 | SRR6257762 | Testis |
| 7955 | SRR6327865 | Liver |
| 7955 | SRR6327866 | Liver |
| 7955 | SRR6327867 | Liver |
| 7955 | SRR6327868 | Liver |
| 7955 | SRR8867570 | Heart |
| 7955 | SRR8867571 | Heart |
| 7955 | SRR8867572 | Heart |
| 7955 | SRS1042402 | ovary |
| 7955 | SRS1042403 | testis |
| 7955 | SRS1095797 | ovary |
| 7955 | SRS1095798 | ovary |
| 7955 | SRS1095799 | ovary |
| 7955 | SRS2867189 | heart |
| 7955 | SRS665978 | brain |
| 7955 | SRS665980 | heart |
| 7955 | SRS665982 | liver |
| 7955 | SRS665983 | kidney |
| 7955 | SRS665988 | testis |
| 7955 | SRS665989 | ovary |
| 7955 | SRS673878 | heart |
| 7955 | SRS673879 | liver |
| 7955 | SRS673881 | brain |
| 7955 | SRS719609 | brain |

| | | |
|---|---|---|
| 7955 | SRS719610 | brain |
| 7955 | SRS719611 | brain |
| 7955 | SRS719615 | heart |
| 7955 | SRS719616 | heart |
| 7955 | SRS719617 | heart |
| 7955 | SRS719621 | kidney |
| 7955 | SRS719622 | kidney |
| 7955 | SRS719623 | kidney |
| 7955 | SRS719624 | liver |
| 7955 | SRS719625 | liver |
| 7955 | SRS719628 | liver |

* SRA ID (https://www.ncbi.nlm.nih.gov/sra)
